# Visualising joint force-velocity properties in musculoskeletal models

**DOI:** 10.1101/2025.06.24.660703

**Authors:** Christopher T. Richards, Tiina Murtola

**Author notes:** Corresponding author: Christopher Richards.

## Abstract

Musculoskeletal modelling opens windows into how muscle properties interact with neural control to govern movement. Though musculoskeletal models produce vast computational data, they lack a visual language which compactly communicates how joint dynamics relate to time-varying muscle activation, force and length change. We developed a novel representation of joint-level force-velocity (joint-FV) properties which shows how agonist and antagonist muscles contribute to overall joint state and its trajectory throughout a movement. Using a model of human goal-directed reaching, we used joint-FV visualisations to discern the salient joint dynamic features across joints and between different reach targets. Regardless of target, we found that the shoulder, elbow and wrist joints traversed a near circular trajectory through joint-FV space when muscle forces were dominant, but trajectories were more complex when joint-interaction forces dominated (i.e. cross-joint forces due to Coriolis, Euler and centrifugal effects). Additionally, we found that co-contraction steepens the instantaneous slope of the instantaneous joint-fv curve causing damping which helps stabilise against small perturbations. We therefore propose that our joint-FV visualisation can be used to explain the intricate features seen in musculoskeletal simulation data to reveal how intrinsic muscle properties govern the behaviour of dynamical systems.

## INTRODUCTION

Musculoskeletal modelling is a powerful tool for understanding how physiological and anatomical properties of muscle not only enable versatile behaviour (Kargo et al., 2002; Kargo & Rome, 2002), but also limit locomotor performance (Clemente et al., 2024). Profound technical advances, such as open-source physics engines (Todorov et al., 2012) and optimisation techniques (De Groote et al. 2016; Dembia et al., 2020), have enabled broad exploration of musculoskeletal biomechanics both in humans and in other animals (Seth et al., 2018). However, the generation of fundamental knowledge from musculoskeletal modelling approaches is not necessarily straightforward given the richly detailed data they produce. Thus, despite the wide availability of modelling tools (Damsgaard et al., 2006; Delp et al., 2007), there remain gaps in our detailed understanding of how muscle dynamics interact with the intricate physics of biomechanical systems.

A primary reason why simulated data from muscle models can be challenging to analyse and visualise is that they churn out multiple streams of time-series data (e.g. force, length, velocity, activation, force-velocity gain). In particular, the question, *What is the functional role of muscle X?* is often difficult to address thoroughly; though computational methods for discerning patterns do exist, visualisation of those patterns is often cumbersome. This is because of the great demands of interpreting large numbers of time-series plots and understanding interactions among them. Nevertheless, visualisation has long been an important tool for communicating key structural properties of time-series data (Muller & Schumann, 2003). Moreover, novel visualisations can further drive generation of knowledge, particularly with dynamic biomechanical systems (e.g. visualisation of skeletal movements using XROMM; Brainerd et al., 2010).

Fortunately, a significant advance came from plotting muscle force against displacement to create a “work loop” figure (Pringle & Tregear, 1969; Josephson, 1985) which collapses muscle dynamics into a trajectory whose shape and direction visually communicates function; i.e. whether the muscle acts like a motor, brake, strut or spring (Full et al., 1998; Ahn & Full, 2002; Dickinson, 2010). To probe further, one can then address the question, *How does the muscle achieve its function?* This is done by observing where the muscle operates on its force-velocity (FV) curve (see Ahn & Full, 2002). For example, during early stance in human running, ankle extensors achieve “strut-like” function due to their operating range on the FV curve during high activation (Arnold et al., 2013). Furthermore, FV effects (in the form of gain functions) can be plotted as time-series to demonstrate the relative impact of activation versus length versus velocity effects on the temporal features of force production during dynamic activity (Clemente & Richards, 2012; Richards & Clemente, 2012). Unfortunately, achieving this level of functional and mechanistic detail requires several plots for each muscle. Hence, these types of zoomed-in analyses of muscle functional mechanics are often restricted to “simple” models (i.e. one or a few muscles; e.g. Curtin et al., 1998; Aerts & Nauwelaerts, 2009; Lichtwark, G. A., & Wilson, 2005; Richards & Sawicki, 2012) whereas a similarly detailed mechanistic analysis for complex musculoskeletal models (e.g. Pandy et al., 1990; Kargo & Rome, 2002; Hutchinson et al., 2015; Arnold et al., 2016; Park et al., 2022) is not always practical, given the number of plots required.

In addition to the data presentation problem, musculoskeletal models have many interacting levels of dynamic behaviour which are difficult to digest. At one level, musculoskeletal models are digital marionettes made of linked segments which can behave counterintuitively and chaotically due to cross-joint interactions (e.g. Hollerbach & Flash, 1982). These multibody systems require extensive nonlinear equations of motion (e.g. Murray et al., 1994) which make intuitive grasp of the physical behaviour challenging. At another level, muscle properties such as force-length-velocity and activation dynamics introduce additional layers of nonlinear behaviour (e.g. Winters, 1990) which might grow even more difficult to comprehend during co-contraction. Hence, a holistic and thorough understanding of musculoskeletal dynamics is profoundly challenging to reach and communicate in a single study.

To help overcome the impracticalities of analysing arbitrarily complex musculoskeletal models, we aimed to consolidate muscle mechanical data into a compact visual format. We introduce a new conceptual framework which visually encapsulates multiple streams of data onto a single plot. We propose a visualisation of “joint force-velocity space” which consolidates muscle data from agonist and antagonist muscles to create a portrait of joint mechanical function. We then use joint-force-velocity plots to unravel how joint-level behaviour is explained by mechanical interactions among antagonistic muscle pairs.

To demonstrate our new framework for visualising muscle mechanical data, we explore the detailed muscle dynamics of a human reaching model (Murtola & Richards, 2023) which encapsulates all the challenges discussed above. We pose three questions: 1) Do the three arm joints function similarly to each other during reaching? 2) Does each joint function similarly across different reaching motions? 3) How does co-contraction influence joint-level force-velocity properties? We found that although generic answers to these questions could be discovered from traditional time-series plots, additional details would have been missed without a visualisation of FV properties at the joint level. Thus, we propose that our novel framework takes one step towards tacking the challenge of understanding muscle dynamics within musculoskeletal models.

## THEORY AND METHODS

### Constructing the joint-level force velocity (joint-FV) space

In a traditional Hill-type muscle model, the familiar force-velocity (FV) relationship is one of the key features which governs the dynamic behaviour of a muscle-mechanical system. Though the force-length (FL) relationship might play an important role in slow movements, for the current study we assume FV effects to be dominant (see Discussion) because of the substantial velocities and time-varying loads during reaching (Murtola & Richards, 2023). In Hill-type models, the FV relationship is drawn as a single curve representing maximum activation such that the normalised muscle force reaches 1 (i.e. maximum force) when velocity is zero (Fig. 1A). Equivalently, the FV relationship can also be understood as a space between two curves where the FV curve defines the ceiling (maximum activation), and the x-axis is the floor. The muscle operates on or between these boundaries. Specifically, when activation is submaximal, the Hill-model FV curve is scaled downwards. Thus, FV space is made from an infinite number of lowercase “fv” curves representing different levels of activation. Consequently, a muscle’s functional state may move through FV space in two ways: 1) by travelling along its current fv curve (constant activation) due to changing load velocity or 2) by moving across fv curves (changing activation). During functional behaviours, a muscle generally moves from one region of the space to another. For example, a muscle which shifts between acceleration (e.g. jumping) and stabilisation (e.g. landing) moves from a region of high power (i.e. ∼1/3 maximum contraction speed) to near isometric. How quickly a muscle’s state can change depends on load as well as the shape of the FV curve and the rate of activation change (as measured by the rising and falling slopes of isometric twitch or tetanic contractions).

**Figure 1.**
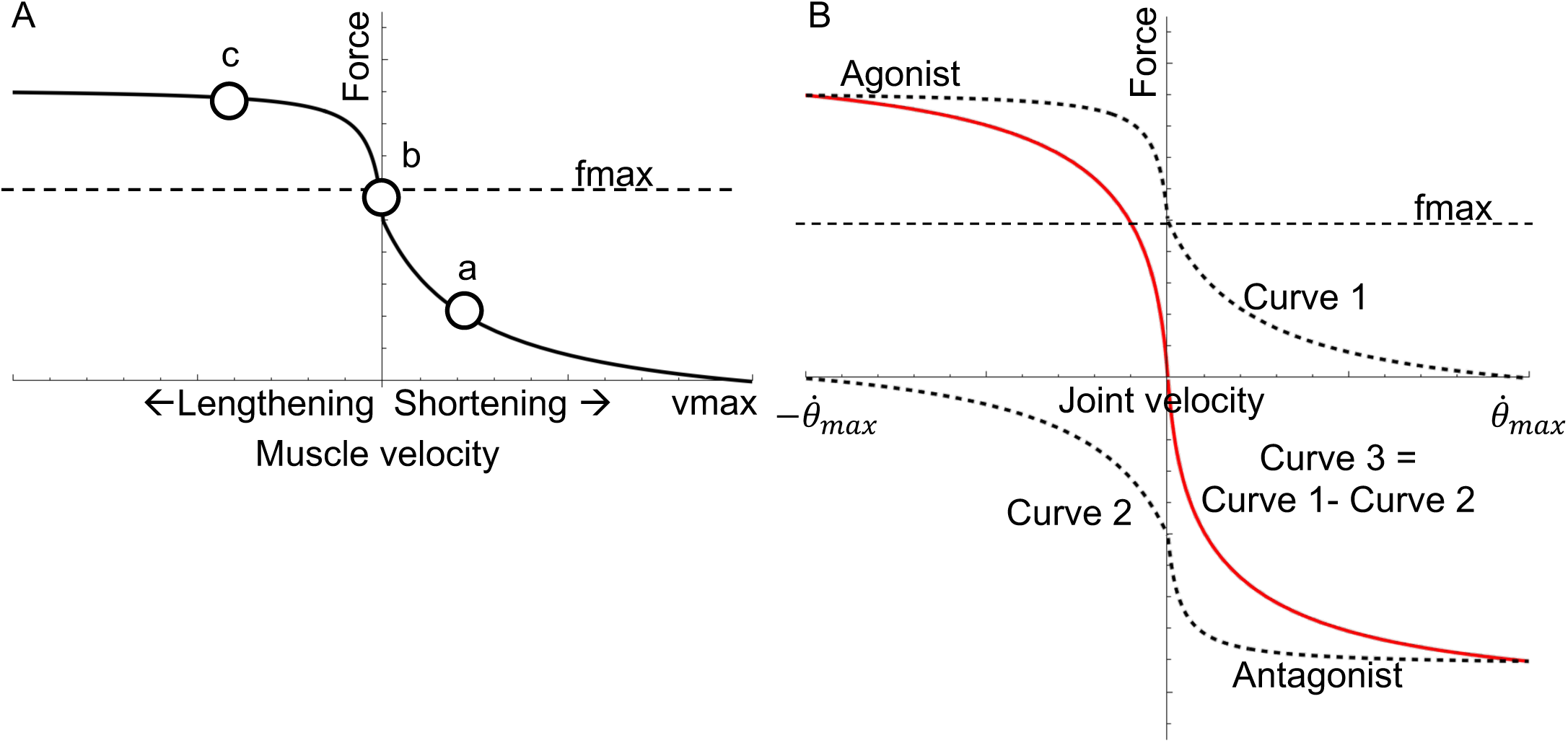
The force-velocity relationship. Muscle force falls hyperbolically with shortening velocity but increases steeply with lengthening (A). Much of dynamic muscle function can be understood by a muscle’s operating point on its force-velocity curve. The diagram shows three hypothetical operating points on the curve (white circles). Point a represents a muscle shortening at ∼1/3 vmax which is optimal for mechanical power (e.g. a muscle at optimal cycling frequency such as in a swimming fish, e.g. Curtin & Woledge, 1988; Rome et al., 1988 or a bird at takeoff, e.g. Askew & Marsh, 2001) versus point b where the muscle operates isometrically (e.g. a leg muscle in a running turkey remains isometric while work is performed by the tendon; Roberts et al., 1997) versus point c where the muscle resists lengthening to absorb energy (e.g. a stabilizing muscle of a running cockroach; Full et al.,1998; Ahn & Full, 2002). Joint-fv relationship (B). The FV curves of the agonist (curve 1, upper, dashed) and antagonist (curve 2, lower, dashed) are identical in joint velocity space, but reflected about the x and y-axes. The agonist curve intercepts the x-axis at +vmax whereas the antagonist intercepts at –vmax. Subtracting curves 1 and 2 gives the joint-fv curve 3 (solid red). Note that the joint-fv curve (curve 3) has local regions of steeper slope than either of its component curves, especially near the origin.

While single-muscle contraction characteristics are governed by its FV relationship, musculoskeletal behaviour depends on how the FV relationships of antagonistic muscle groups interact at the joint level. We therefore introduce the concept of a “joint-FV space” which describes all combinations of joint net force and joint velocity that are possible given the FV relationships of the muscle groups. In the general case with multiple mono- and biarticular muscles crossing a single joint, the shape of the joint-FV space is complicated. However, in the present work, we focus on the simplest case of a symmetric monoarticular joint with a single pair of identical antagonistic muscles (more complex configurations are possible, but outside of the current scope; see Discussion). In this simple case, the joint-FV space can be constructed by considering how the velocities of the two muscles are linked through the joint angular velocity. Consequently, the boundaries of possible joint-FV states are defined by the FV curves of the two muscles where the antagonist curve has been reflected about both the x- and y-axis (because if one muscle shortens, the other must lengthen at the same speed, and the forces/torques of the two muscles act in opposite joint directions; Fig. 1B). The position of a muscle pair in this joint-FV space depends on the activation levels of the two muscles. Specifically, these activation levels define the current joint-fv curve (analogous to the single-muscle fv curve) and a local joint-fv space that spans the family of joint-fv curves between the two single-muscle fv curves (i.e. shaded region in Fig. 2A). The joint-fv curve represents all joint states accessible without changing activation levels, while the joint-fv space gives a visual representation of how each muscle contributes to the joint-fv curve. The morphing of joint-fv space creates a rich variety of possible trajectories through the joint-FV space as activation levels dynamically change during complex behaviours. Notably, when co-contraction occurs, the joint-fv curve differs in shape from the single-muscle fv curve, particularly near the y-axis where co-contraction causes a steeper slope.

**Figure 2.**
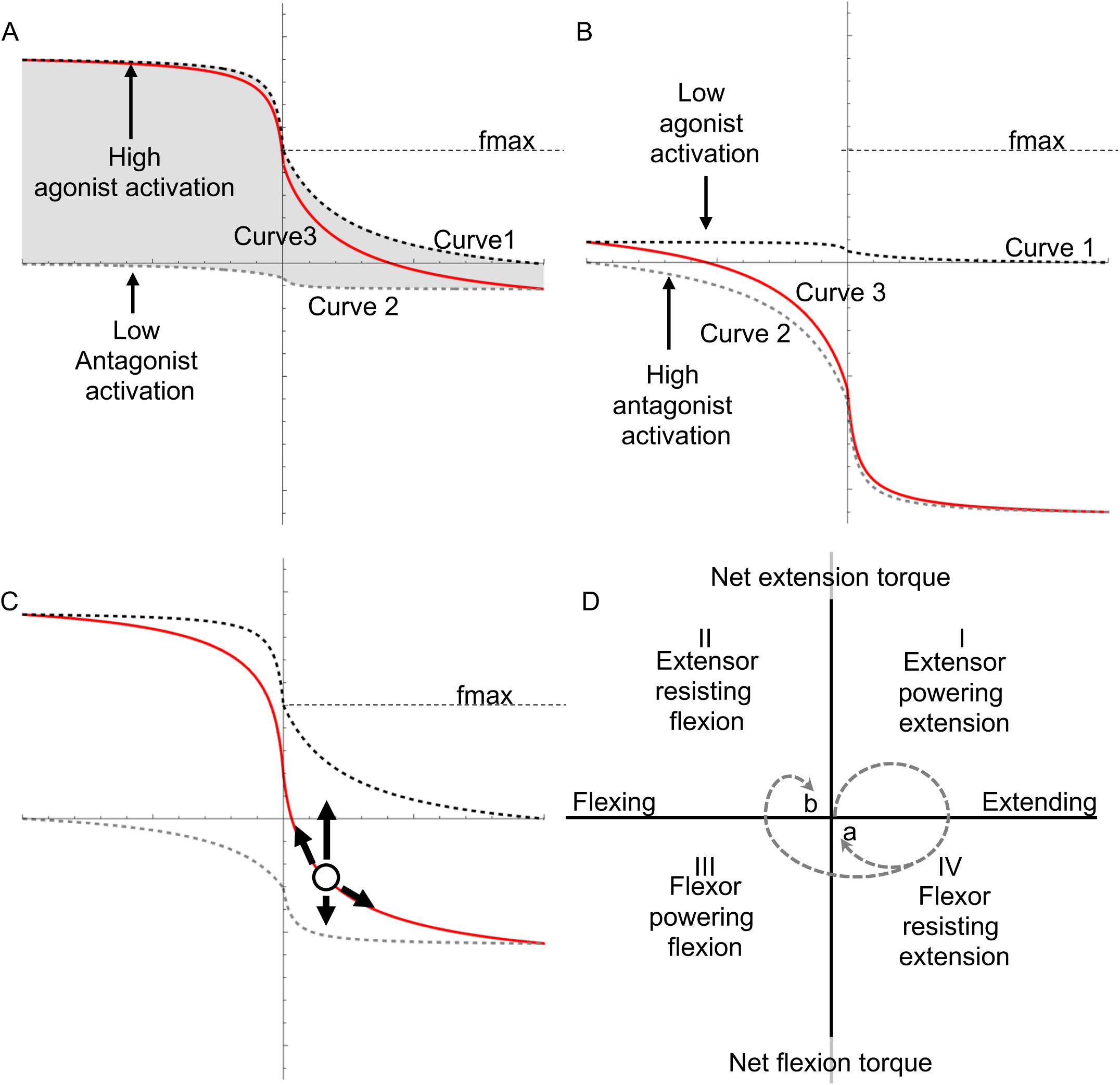
The joint force-velocity (joint-fv) relationship. (A) Curves 1 and 2 represent the upper and lower bounds of the joint-fv space, depending on the respective levels of agonist and antagonist activation. For any agonist/antagonist activation level, the area between curves 1 and 2 (shaded region) represents the “joint-fv space”. At a moment in time, the relative agonist/antagonist activation levels determine the joint’s instantaneous fv curve (red; curve 3). In this example, agonist activation is near maximum and agonist activation is near 0; curves 1 & 3 nearly overlap one another (they will entirely overlap if antagonist activation is 0). (B) In this reciprocal example, agonist activation is very low (compared to the antagonist), hence curves 2 & 3 nearly overlap. (C) This example shows maximum agonist activation, but 50% antagonist activation. The joint operating point (white circle) always falls on its instantaneous joint-fv curve. This point may move either up or down, due to changes in either muscle’s activation, or the point may move along the curve due to changes in mechanical loading. (D) The four quadrants of joint FV space indicate the mechanical function of the joint (e.g. with respect to flexion or extension action). In an ideal reach where the arm produces a monotonic movement precisely to the target, a joint would start from rest at the origin then move along a circular path (dashed path a). More typically, a joint trajectory has an additional homing-in phase (path b). In any case, the movement of the joint through quadrants reflects the time-varying mechanical function of the joint (see text). Note this example shows an extensor as the agonist (e.g. elbow, wrist), however that choice is arbitrary; for the shoulder, the flexor would be the agonist and the flexion/extension labels would swap (see text).

The FV calculations are as follows. The agonist muscle FV curve is converted to joint angular velocity from the FV function used previously (Murtola & Richards, 2023).

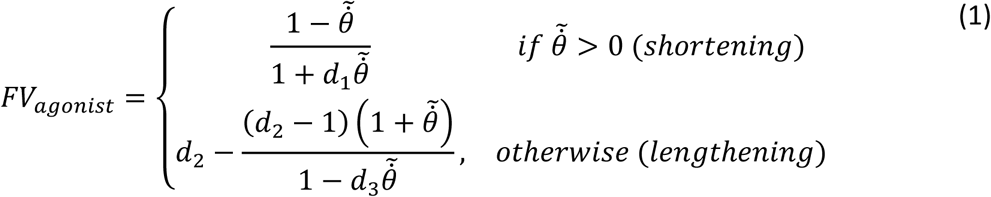

Where *d1*, *d2*, *d3* are shape constants (*d1*=4, *d2*=1.8, *d3*=30.24 in the current study; Murtola & Richards, 2023) and 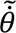 is the normalised joint rotational velocity which is positive for shortening.

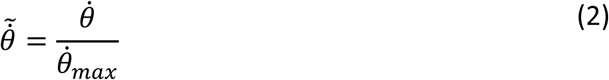

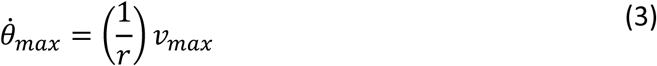

where *vmax* is the maximum muscle shortening velocity (in m/s) and 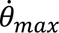 is the maximum joint velocity (in rad/s), given the moment arm, *r*.

The antagonist FV curve is defined as above, except with all velocity terms multiplied by -1 in Eq. 1. In reality, moment arms may be time-varying and differ between agonist and antagonist. For simplicity, and to match our simulations, we assume agonist and antagonist moment arms are equal and constant for the current study, however the framework can be expanded in the future to allow for these complexities (see Discussion).

The joint-fv curve (at one instant in time) is as follows.

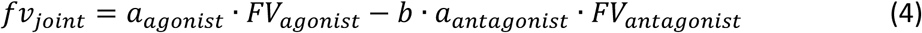

where *aagonist* and *aantagonist* are the activation levels of the agonist (i.e. the dominant muscle) and antagonist, and *b* is a weighting factor, *b = fmaxantagonist/fmaxagonist* to account for different muscle strengths. The joint-fv space is defined as all the joint-fv curves (Eq. 4) that correspond to agonist activations between 0 and current *aagonist* and antagonist activations between 0 and current *aantagonist*. Note that the independent activation levels *aagonist* and *aantagonist* can equivalently be expressed using coactivation, which describes the level of activation simultaneously present in both muscles, and net activation, which describes how much excess activation the agonist has relative to the antagonist: *anet = aagonist – b*aantagonist*.

To generalise Eq. 4 to a variety of tasks, we redefine agonist/antagonist in a way that is independent of specific anatomical movement directions, such as flexion/extension. As the mechanical function of a joint is indifferent to anatomical orientation, we can freely choose which muscles to label as agonists based on the mechanical task. For some tasks, this selection can easily be made to align with anatomical convention, such as during a jump requiring leg extension, where all extensors would be called agonists and flexors would be antagonists. However, reaching is less straightforward because some joints may move in different directions (e.g. elbow extends whilst shoulder flexes) and the direction of joint movement may change during a reach (e.g. during homing-in) causing muscle roles to alter between powering movement, breaking, and stabilising. Furthermore, this pattern may change depending on target location. To address this problem, we define an agonist as the dominant muscle which produces the power to accelerate a load during the first 0.05s of the reach. Furthermore, we define the FV co-ordinate system so that the agonist FV curve is plotted with its shortening portion in the first quadrant of cartesian coordinates whereas the shortening portion of the antagonist appears in the III^rd^ quadrant (Fig. 2D).

### Visualising the joint-fv curve and space

Because of the asymmetry of the FV function, the shape of joint-fv space depends on the relative activations of agonist versus antagonist. At one extreme, the agonist contracts while the antagonist is inactive causing the joint to operate on the ceiling of the space (on curve 1, Fig. 1B). Towards the other extreme, the agonist activation level is low whereas the antagonist activation is high such that the joint operates on the floor of joint-fv space (on curve 2, Fig. 1B). For any other activation values, the joint operates in the area between the ceiling and floor along an intermediate joint-fv curve (curve 3, Fig. 1B; analogous to the fv curves in the single-muscle example above). As the relative activation levels change, the joint-fv space morphs as the space between curves shrinks or expands to represent how each muscle contributes to the joint-fv curve (compare Fig. 2A vs. 2B). For example, low antagonist activation gives rise to a joint-fv space that is mainly above the x-axis while a joint-fv space with similar areas above and below the x-axis indicates a high level of coactivation. Similar to how a single muscle always operates at a point along an fv curve, the joint always operates at a point in FV space along a joint-fv curve (Fig 2C).

### Functional interpretation of trajectories through joint-FV space

The current representation of joint-fv space is analogous to the “work loop” concept of muscle contraction (Pringle & Tregear, 1969; Josephson, 1985). A work loop plot visually communicates muscle function because the work loop shape and direction indicate whether a muscle is producing mechanical work or behaving like strut, brake or spring (see Ahn & Full, 2002). Analogously, the current representation of the joint-fv space can visually communicate joint function through the lens of physiological limitations of activation and FV properties. Specifically, the joint-FV coordinate system can be subdivided into four cartesian quadrants (Fig 2D). For illustrative purposes, we describe the quadrants in terms of flexion and extension, as follows. If the agonist is an extensor and the antagonist is a flexor, quadrant I represents extension powered by extensors. Quadrant IV represents extension but resisted by co-contracting flexors. Quadrant III represents flexion powered by flexors. Finally, quadrant II represents flexion but resisted by co-contracting extensors. Importantly, all muscles can be active in all quadrants, but extensors dominate in quadrants I and II whereas flexors dominate in III and IV. When the joint operates along the y-axis, both muscles are isometric whilst the joint is stationary. The x-axis represents the balance of both muscles meaning no net torque on the joint. For example, the elbow of a jumping frog would likely operate mainly in quadrant I during launch, but in quadrant II during landing. Following from our current definition of agonist and antagonist muscles, the co-ordinate system (Fig. 2D) can be oriented to match the behaviour of interest. For example, a predominantly flexion-powered movement such as a “biceps curl” exercise would have flexors occupy quadrants I and II (versus extensors in III and IV).

The dynamics of joint function can be studied by observing the trajectory of movement through joint-FV space. Similar to the single muscle example, a joint’s operating point can move anywhere in the available space either by moving up and down due to relative changes in agonist/antagonist activation or left and right along the current joint-fv curve due to changes in loading. Specifically, a joint’s operating point in FV space can move straight up as rapidly as the agonist activates (and the antagonist deactivates). However, a joint’s motion along the joint-fv curve is only limited by how rapidly the load can accelerate. For example, a joint working isometrically against a latched load will be on the positive y-axis; once the latch releases, the joint operating point will travel rapidly to the right along the joint-fv curve.

### Reaching simulations

To demonstrate the joint-FV concept, we used a previously published musculoskeletal model of human goal directed reaching (Murtola & Richards, 2023). Briefly, the model is based on simplified human anatomy and inertial properties, and it is restricted to the horizontal plane (Fig. 3A). The movements of the hinge joints at the shoulder, elbow and wrist, are each driven by a pair of monoarticular Hill-type muscles. The arm is guided towards the target by muscle excitation signals computed by a PD controller following a pre-defined trajectory. All muscle properties are identical to our previous study; however, the parameters of the PD controller were retuned for rapid reaching (i.e. to reach the target within ∼0.5s).

**Figure 3.**
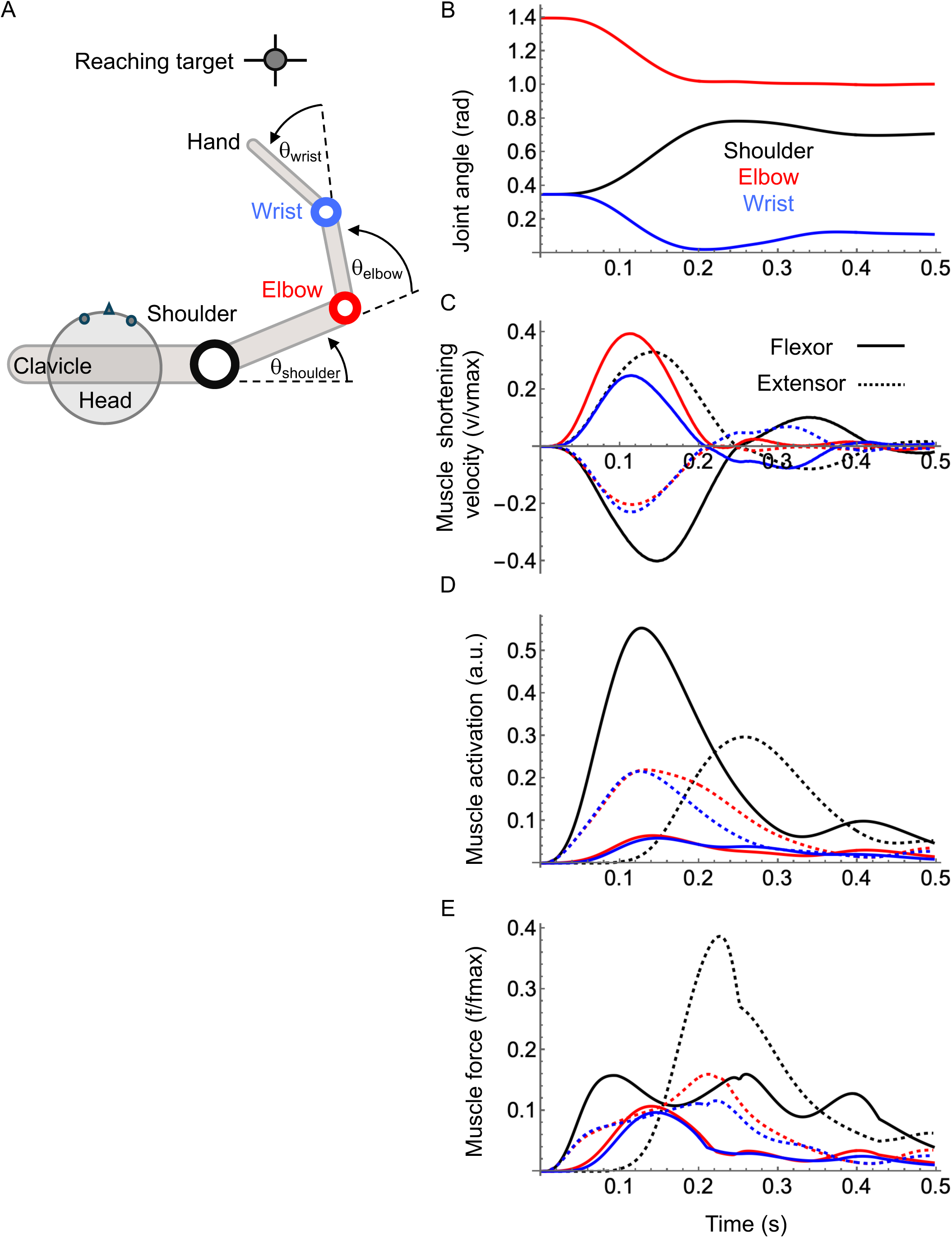
Musculoskeletal simulation of human goal-directed reaching. (A) A schematic of the horizontal reaching model (see Murtola & Richards, 2023) consists of an antagonistic pair of muscles about each of its three joints (shoulder, elbow, wrist). Time series data from a simulated reach showing joint angle (B), shortening velocity (C), activation (D) and normalised force (E) for muscles at the shoulder (black), elbow (red) and wrist (blue). Flexors are shown with solid lines whereas extensors are dashed.

To represent reaching over a range of conditions, we selected four conditions to compare: a) forward target; b) sideways (ipsilateral) reach; c) cross-body (contralateral) reach; d) forward target, identical to (a), but with an external perturbation. The perturbations were modelled by injecting a 100N point force (∼1/20^th^ the force of a punch; Adamec et al., 2021) to the hand segment delivered at 90° anti-clockwise to the instantaneous hand velocity vector, ∼150ms after the start of the reach. We were careful to use a perturbation small enough to ensure the perturbed muscle speeds did not exceed *vmax* to avoid violating the assumptions of the Hill-type muscle model. Our code allows for arbitrary control of perturbation magnitude, direction and timing; however, a full sweep of perturbation parameters is beyond the current scope (see Discussion).

## RESULTS

### The muscular dynamics of forward reaching

The forward reach is a simple countermovement between the upper arm and lower arm where the shoulder flexes whilst the elbow and wrist extend smoothly towards the target (Fig. 3B; SI movie 1). The movement has two phases: 1) the primary phase where the arm gets close to the target followed by 2) a final correcting phase where hand homes-in on the target. This is most clearly visible in the “two hump” pattern of muscle shortening velocities (Fig. 3C). The muscle activation patterns (Fig. 3D), however, are more complex and marked by strong co-activation and therefore co-contraction of flexors/extensors at all joints (Fig. 3E). The intricate pattern of muscle forces stems from the inherent complexity of multibody systems where segment rotational velocities and accelerations are mutually dependent via nonlinear relationships. That is, some features of any given force profile are muscle contractions aimed at accelerating its respective segment, whereas other features work to counteract interaction forces between segments. Regardless of the complexity of force patterns, the co-contraction is a salient feature across a broad range of reaching movements due to the intrinsic delays of realistic twitch and tetanic activation profiles, as represented by the 3^rd^ order dynamics used in our model (Murtola & Richards, 2023).

### Joint-FV dynamics of the shoulder, elbow and wrist during forward reaching

Because the simulations start at rest, all joint trajectories begin at the origin of the joint-FV coordinate system. For the shoulder, the dominant action is flexion; therefore, the flexor muscle (agonist) drives the early part of the reach (Fig. 4A; SI movie 3). As agonist activation rises, the joint-fv space expands and the joint travels along a parabolic trajectory until net activation reaches a peak (Fig. 4B) then subsequently falls due to rising antagonist (extensor) activation. To slow the arm and minimise overshooting the target, the extensor then resists the flexion (Fig. 4C). In the final stages, the activation of both muscles falls, shrinking the size of the joint-fv space as the joint moves from flexion to extension to correct overshoot and home in on the target (Fig. 4D,E,F). Overall, the trajectory is a clockwise path through FV space (similar to “path a” in Fig. 2D).

**Figure 4.**
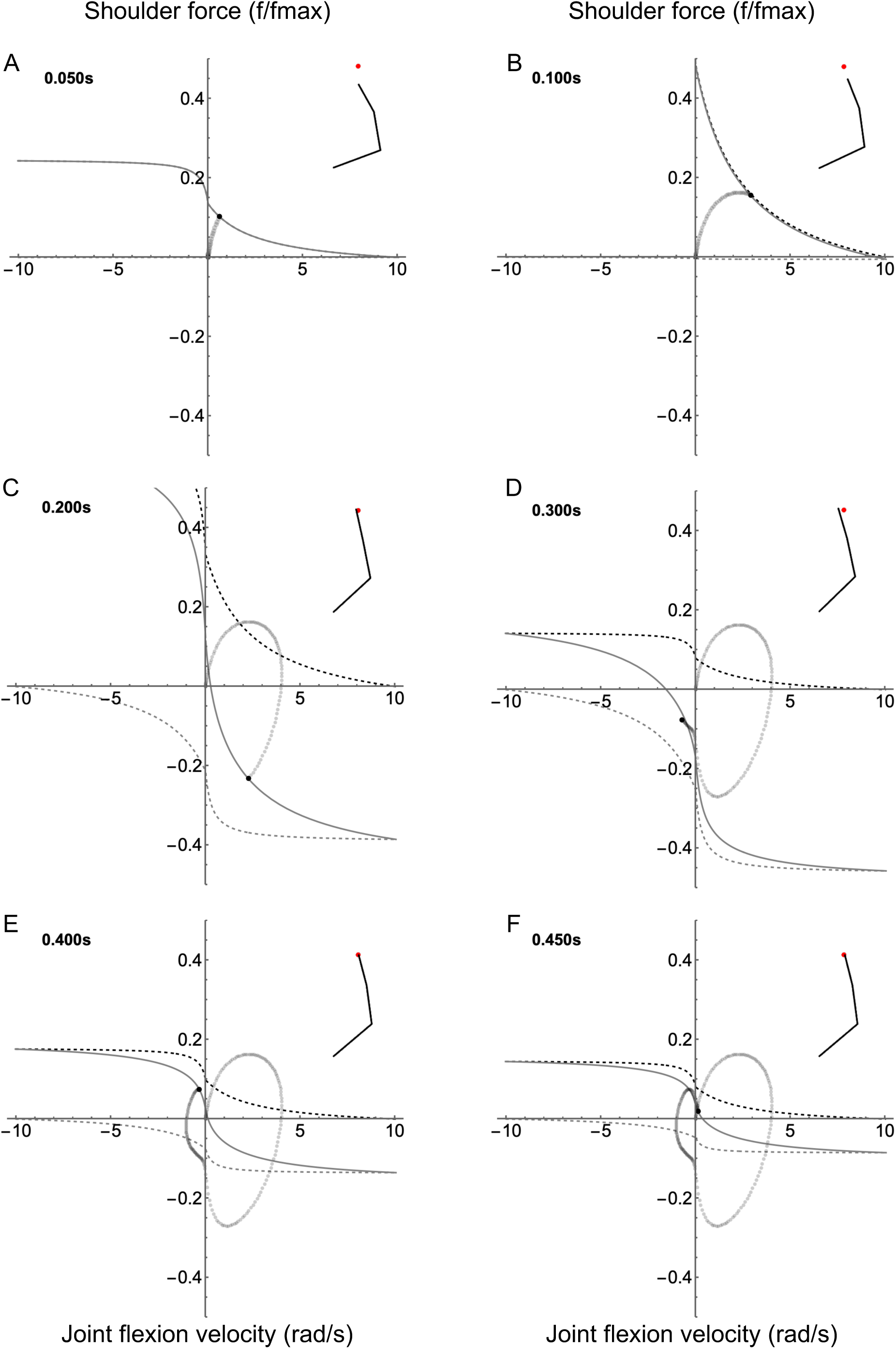
Joint FV space animation for the shoulder joint. FV curves are shown for 6 time frames throughout the reach (A-F). For the shoulder, the flexor is defined as the agonist, therefore positive force causes flexion. The inset shows position of the right arm relative to the target (red dot). The joint state trajectory is shown as a trail of gray dots, with the current state indicated by the black dot. The solid black line is the current joint-fv curve, and upper and lower limits of the joint-fv space are indicated by the dashed black and gray lines, respectively.

Similar to the shoulder, the elbow joint-fv space begins small (Fig. 5A; SI movie 3) then expands to a peak (Fig. 5B, C) then falls throughout the reach (Fig. 5D,E,F) similar to the fall in activation of both muscles observed in the shoulder. Unlike the shoulder, the dominant action of the elbow is extension. Though the early phase of the reach is similar to the shoulder, the elbow remains in quadrant I for the majority of the reach indicating that its main role is to produce power to move the arm towards the target. In the final homing-in stages of the reach (Fig. 5D, E, F) the joint remains isometric as it stabilises the arm about the target position.

**Figure 5.**
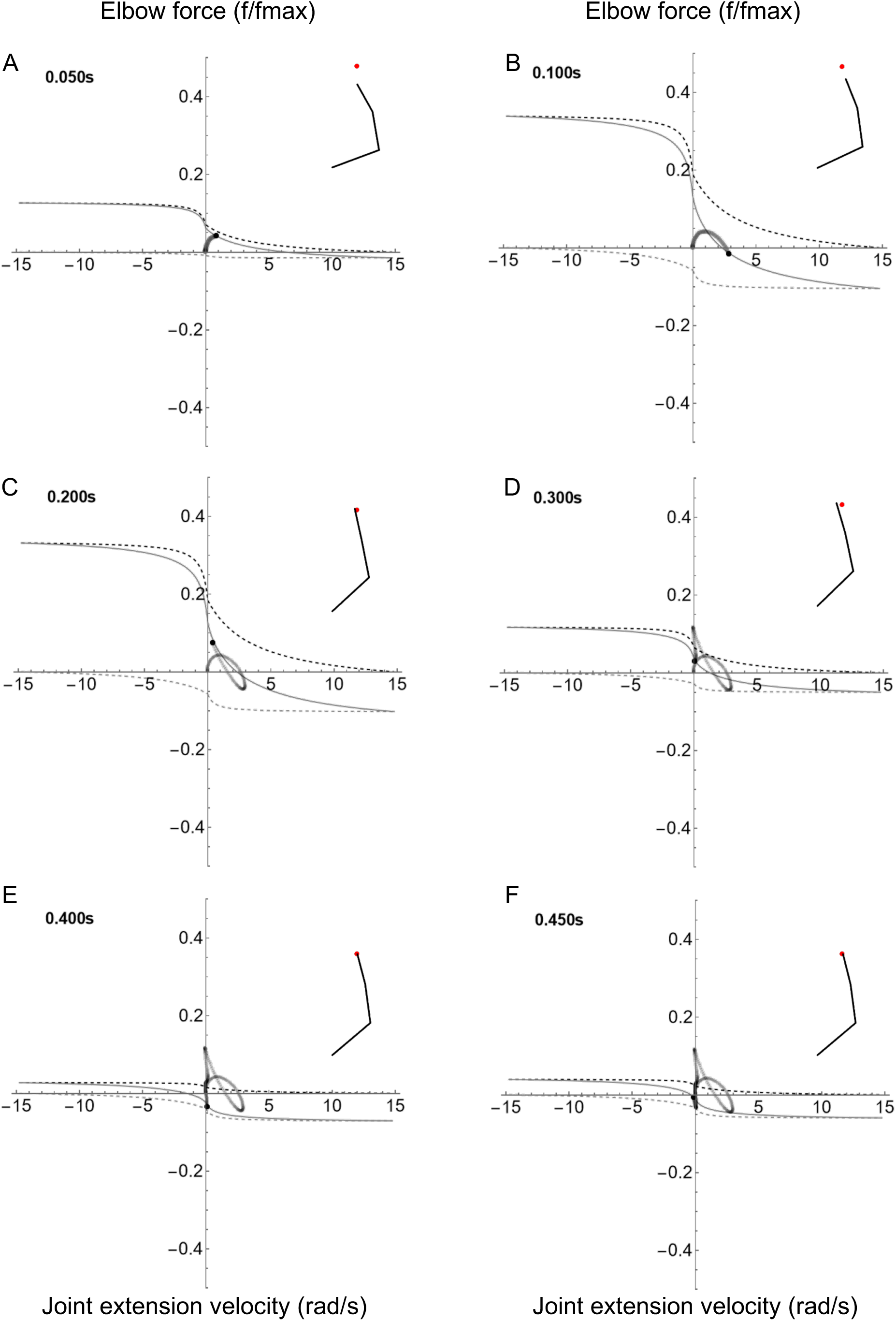
Joint FV space animation for the elbow joint. FV curves are shown for 6 time frames throughout the reach (A-F). For the elbow, the extensor is defined as the agonist, therefore positive force causes extension. The inset shows position of the right arm relative to the target (red dot). The joint state trajectory is shown as a trail of gray dots, with the current state indicated by the black dot. The solid black line is the current joint-fv curve, and upper and lower limits of the joint-fv space are indicated by the dashed black and gray lines, respectively.

The wrist resembles the other joints in terms of the rise and fall of the size of the joint-fv space (Fig 6; SI movie 3). Like the elbow, the dominant movement of the wrist is extension. Additionally, the wrist functions like the elbow regarding its trajectory through the joint-FV space.

**Figure 6.**
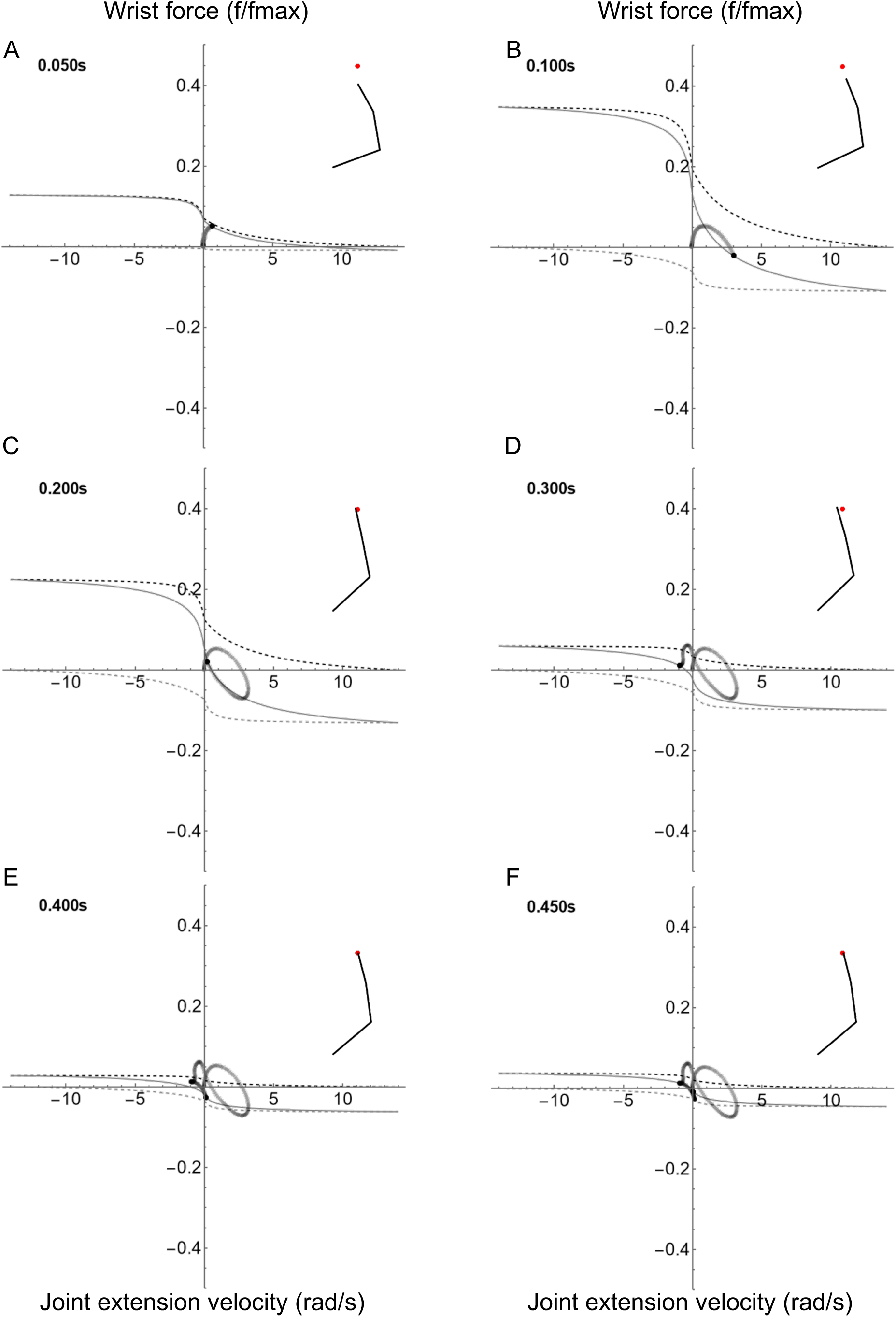
Joint FV space animation for the wrist joint. FV curves are shown for 6 time frames throughout the reach (A-F). For the wrist, the extensor is defined as the agonist, therefore positive force causes extension. The inset shows position of the right arm relative to the target (red dot). The joint state trajectory is shown as a trail of gray dots, with the current state indicated by the black dot. The solid black line is the current joint-fv curve, and upper and lower limits of the joint-fv space are indicated by the dashed black and gray lines, respectively.

### Comparing joint-FV dynamics of forward vs sideways vs cross-body reaching

Comparison of forward, sideways, and cross-body reaching showed broadly similar joint-fv trajectories, however some unique features emerged depending on joint and reach type (Fig. 7; SI movies 3, 4, 5). Overall, we identified three qualitative categories of joint-fv trajectory: 1) the circular-like path where the quadrants are traversed in sequence with a primary phase in quadrants I and IV and an overshoot-corrective phase in quadrants III and II (Fig. 7A, B, C), 2) the “tied loop” path where the loop starts to cycle through quadrants I and IV as above, but returns to the first quadrant before homing in, thus creating a “tie” at the beginning/end of the loop (Fig. 7D, E, G, H) and 3) the “fish-like” path where the early reach is similar to other paths, but the latter half of the reach has an elongated “tail” in quadrants II and III as the joint moves strongly in the opposite direction (Fig. 7F, I). Type 1) is exemplified by the shoulder joint which is the most strikingly similar across reaches; its circular clockwise path indicates that joint movement is mostly driven by the shoulder muscles and characterised by agonist-powered acceleration in one direction, followed by antagonist braking and then a corrective movement in the opposite direction. This corrective movement is the smaller secondary loop (“ear”) in quadrants II and III (Fig. 7A, B, C). In contrast, the elbow and wrist joints all showed more complex joint-fv trajectories including anti-clockwise loops or isometric contractions which are due to complex dynamics of inter-segment effects rather than contraction of the muscles moving the joint (see Discussion).

**Figure 7.**
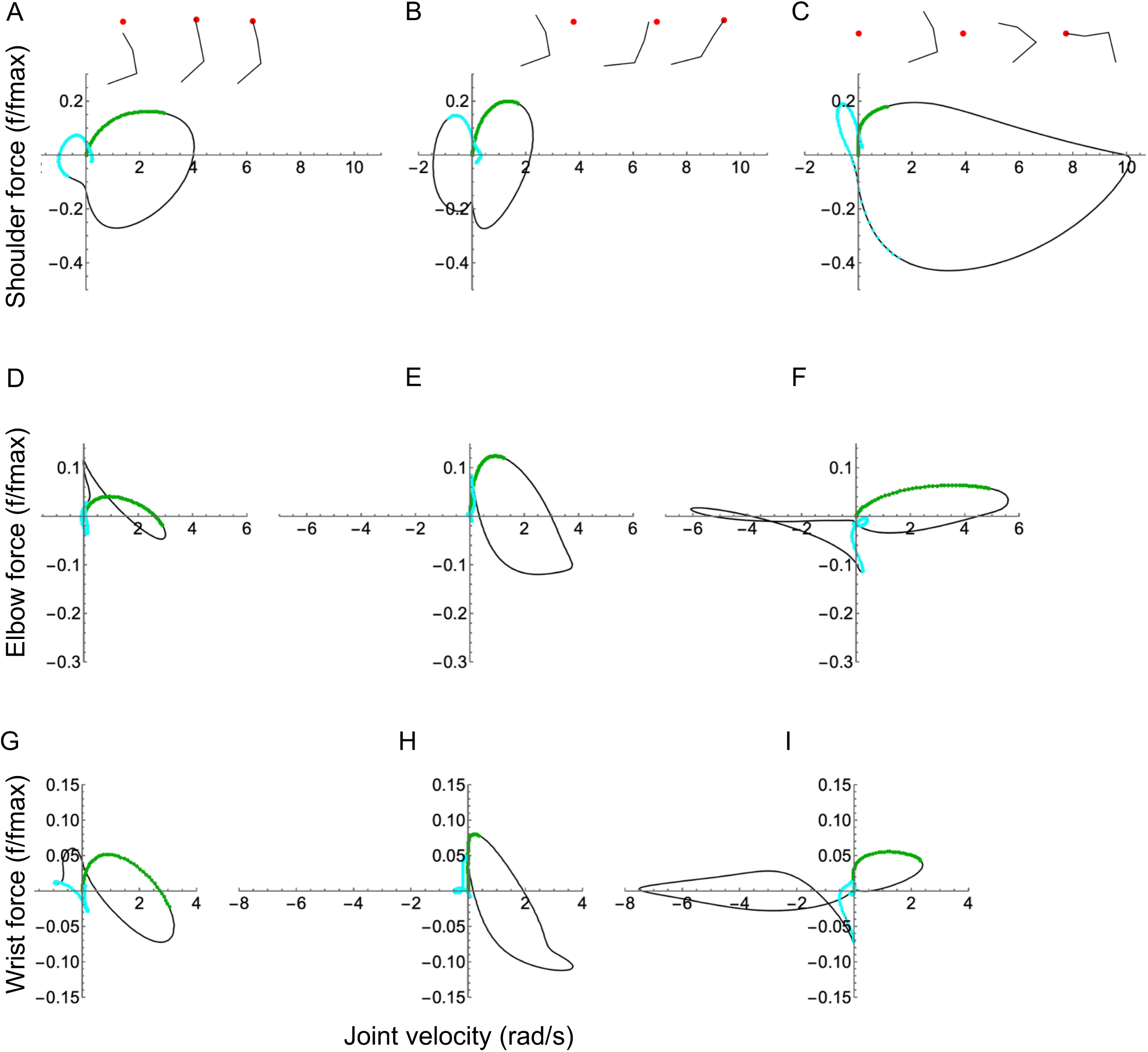
Joint-fv trajectories for the shoulder (top), elbow (middle) and wrist (bottom). Left column shows the forward reach (A,D,G), middle column is the sideways reach (B,E,H) and the right column is the cross-body reach (C,F,I). Top insets show stick animation frames of the right arm and target (red dot). The thick green curve segment highlights the first 0.1s of simulations and the thick cyan highlights the final 0.2s of simulations.

### The effects of a small mechanical perturbation on joint-FV dynamics

Reaching kinematics were slightly altered in the presence of a small mechanical perturbation, however the impact was not strong enough to affect the reaching outcome (SI movie 2). The effects of perturbation varied depending on the joint, however we show the effects at the elbow because they are most pronounced and best illustrate the utility of the joint-FV visualisation. The perturbation caused a minor dip in the elbow torque followed by a correction to allow the arm to reach the target successfully (Fig. 8A vs 8B; SI movie 6). There were two phases of the correction stage. First, a rapid rebound phase returned the torque to its planned trajectory. This response can be seen in the extremely rapid movement of the joint’s position in joint-FV space. Second, a slower adjustment (mediated by the controller) modulated muscle activation to produce torque to compensate for the increased error during the perturbation. Differences between control and perturbation are most obvious when viewing the torque rate (i.e. its time derivative). Most strikingly, a rapid rebound correction occurs despite only modest changes in activation rate and force-length gain (additionally, moment arms are nearly constant). This suggests that FV effects play a large role in the most rapid part of the perturbation response. Specifically, the co-contracting muscles produced a joint-fv curve with steeper slope than either muscle acting alone (Fig. 9A), possibly enabling the joint to respond more rapidly than a “controlled” response.

**Figure 8.**
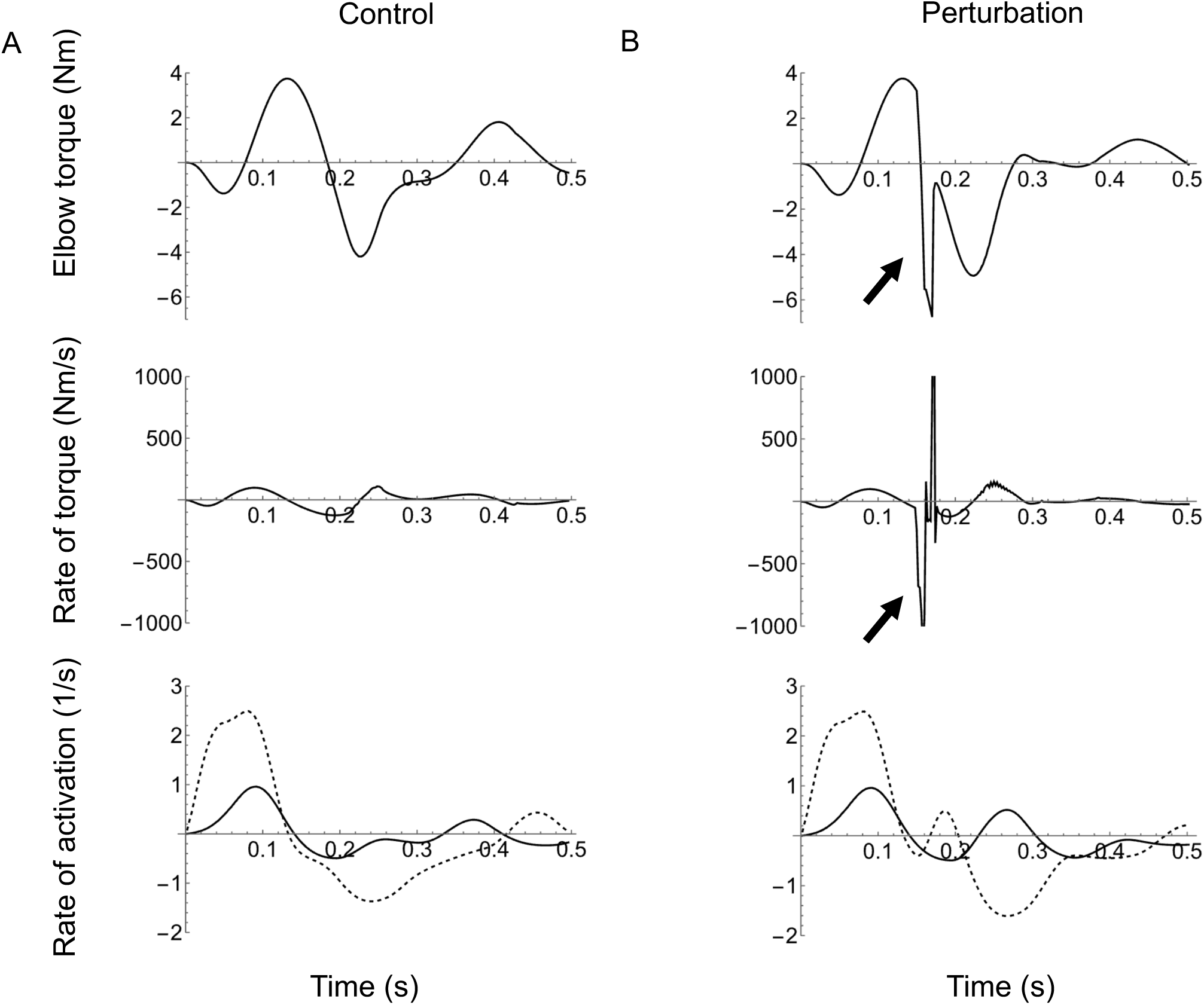
The effect of perturbation. Elbow joint torque, rate of torque (Black) as well as the rate activation (red) for the flexor (solid) and extensor (dashed) for control (A) versus perturbation (B) conditions. The arrows indicate the spike in torque and torque rate due to the perturbation. Note how during the perturbation response, the transient rate of torque exceeds the single-muscle limit of activation rates and likely emerges from extremely rapid movement along the FV curve (see text).

**Figure 9.**
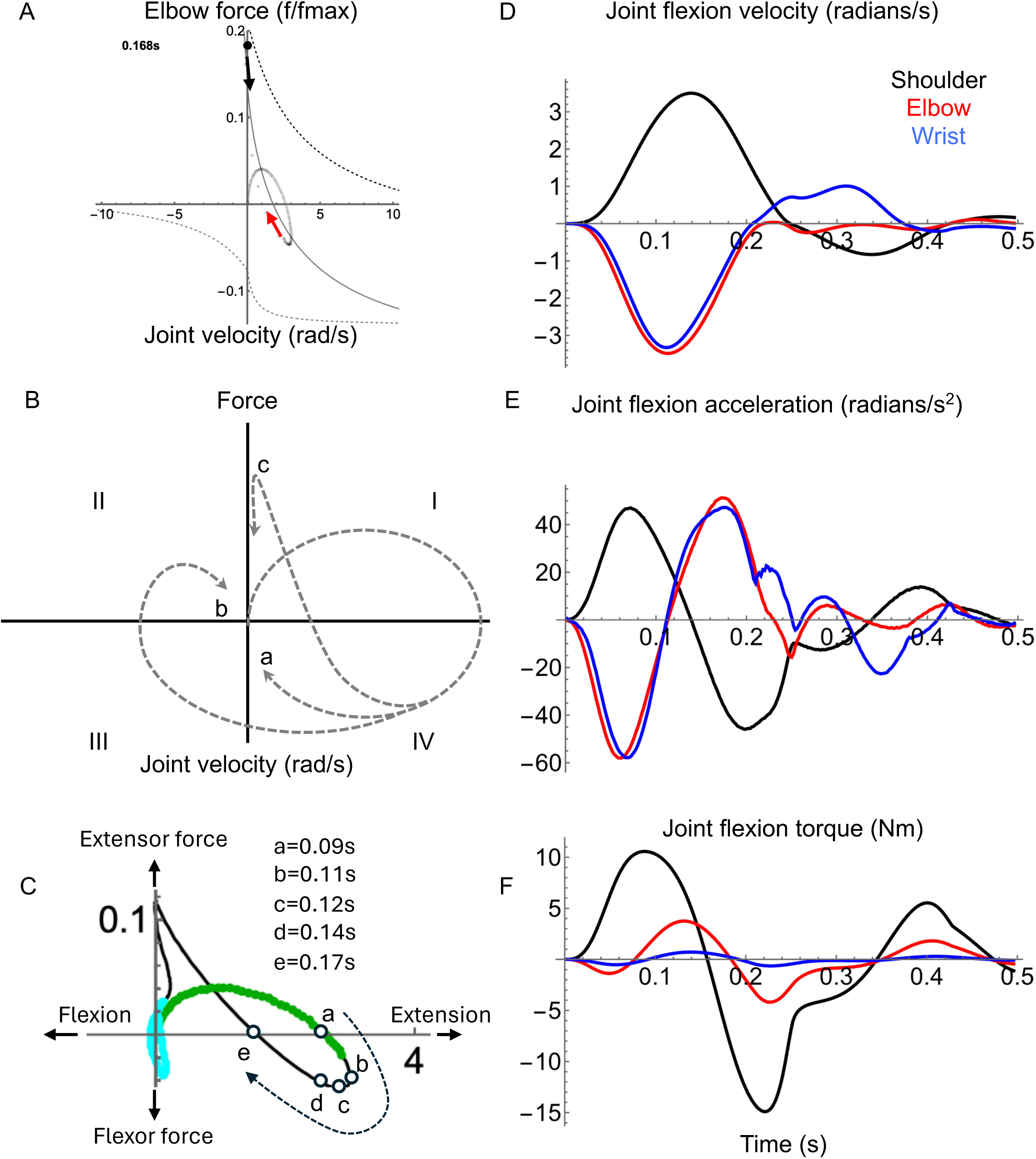
Effects of external forces. (A) A snapshot is shown from the elbow joint-fv trajectory at the instant just following the perturbation (same axes as in Fig. 5). At the perturbation, the joint is pushed upwards/leftwards along its joint-fv curve (red arrow). After the perturbation, the joint rebounds rapidly back down the region of the joint-fv curve where the slope (therefore mechanical damping) is highest (black arrow). The black dot is the current joint state. (B) Idealised joint-fv trajectories for non-perturbed reaching (as in Fig. 2) showing (a) a perfectly planned single-joint path, (b) a feedback-corrected path similar to shoulder trajectories, and (c) a path with an “external force” due to multi-joint interaction torques similar to elbow and wrist trajectories (see text). (C) The elbow trajectory (axes and colours as in Fig. 7A) with white circles showing trajectory time points of interest (a-e mark time points of 0.09, 0.11, 0.12, 0.14 and 0.17s) to aid the description in the text (see Discussion). (D) Angular velocity, (E) acceleration and (F) net joint torque for shoulder (black), elbow (red) and wrist (blue) joints during the forward reach (unperturbed). Note how the shoulder torque roughly follows joint acceleration. In contrast, elbow and wrist acceleration are dominated by cross-joint interaction torques and, consequently, the joint accelerations deviate from the net muscle torques, causing the joint trajectory to follow path c rather than a or b.

## DISCUSSION

We aimed to develop a visual representation of joint mechanics in terms of joint-level force-velocity (joint-FV) properties. To demonstrate this novel representation, we ran simulations with a previously published reaching model (Murtola & Richards, 2023) and plotted the data in the joint-FV coordinate system. For ease of discussion, we adopt the useful abstraction that joints themselves “have actions”; i.e. a joint can be described as producing torque, work or power (see Roberts & Belliveau, 2005) with the implicit understanding that a “joint” in the musculoskeletal modelling sense is not an active tissue structure, but represents the net behaviour of all muscle and tendon actions crossing it.

There are three main results, corresponding to the questions posed in the introduction: 1) For any of the three representative reaches, the shoulder, elbow and wrist functioned similarly to each other in terms of gross joint kinetics. During the first phase, agonist muscles accelerated the arm towards the target. In the second phase, the antagonist muscles engaged to slow the limb, to minimise overshoot and to home-in on the target (Figs. 4, 5, 6). Beyond these gross similarities, the shoulder functioned differently to either elbow or wrist. Specifically, the shoulder acted more like an isolated joint whose motion corresponds tightly to the net torque produced by its muscles. This was not the case for the elbow or wrist whose movement patterns were strongly influenced by shoulder movement in addition to local muscle torque (see below). 2) Joints functioned differently for the three reach targets. However, the qualitative differences were greater among joints than between reaching targets, especially for the shoulder compared to the elbow and wrist (Fig. 7). 3) Co-contraction caused a steepening of the instantaneous joint-fv curve (Fig. 9A) enabling a damping-mediated rebound from small perturbations (Fig. 8B).

### The joint-fv trajectory is a simple and concise portrait of joint mechanical function

The joint-fv trajectory is extremely rich with information. This information can be read at two levels. At the most basic level, the joint-fv trajectory can be read as a visual description of joint behaviour (regardless of the joint-fv curve shape); it is a portrait of joint kinetic function throughout a movement. For example, during the sideways reach, the shoulder action can be divided into four phases with extension during the first two phases (right quadrants) then flexion during the final two (left quadrants). Over this motion pattern, the shoulder undergoes an alternating cycle of generating mechanical power (positive torque for acceleration; quadrant I) to absorbing mechanical power (negative torque for deceleration; quadrant IV) to generating and then absorbing power again (quadrants III and II, respectively; Fig. 7B). The first and second half of the reach correspond roughly to the submovements of reaching (see Meyer et al., 1988) where the arm initially gets close to the target (primary submovement) then performs corrections and homing-in movements (secondary submovements).

Additionally, we view the joint-fv trajectory analogously to a work loop plot (Pringle & Tregear, 1969; Josephson, 1985). In this view, the joint-fv trajectory portrays the net muscle function about a joint and indicates how dominant muscles’ mechanical energy contributes towards net joint function (i.e. behaving like what is traditionally called a motor versus a strut versus a brake; see Dickinson, 2010). For example, the shoulder acts like a flexion motor during the early phase of reaches (until ∼100 ms) driven by the dominant motor behaviour of the flexor. During the middle phase of reaches, the shoulder acts as a brake to slow the joint (action dominated by breaking by the extensor), then later reverses to an extension motor to correct the overshoot of shoulder flexion. Likewise, the elbow and wrist also begin as extensor motors, but the elbow differs dramatically from shoulder function during the homing-in phase. Rather than reversing direction, the elbow produces an early braking torque, then functions isometrically (like a strut; see Fig.5 C-F where the joint-fv trajectory moves vertically along the y-axis). The wrist shares behaviour of both elbow and shoulder, showing early braking function (elbow-like) followed by late-stage extension-flexion reversal (shoulder-like).

At a second level, we go beyond a basic kinetic or energetic description by considering the underlying muscle force-velocity dynamics to explain *why* the joint behaves as observed. Although the traditional motor-brake metaphors (Dickinson, 2010) are useful at the single-muscle level, joint level function requires more nuanced language to capture the joint’s mechanical state. For example, all joints might be traditionally called “motors” during the first ∼100ms of reaches (clockwise trajectory in quadrant I), but the shoulder generates movement mainly through pure agonist action while the elbow and wrist modulate mechanical power production through co-contraction, as visualised by the size and location of the joint-fv space. This co-contraction makes the wrist and elbow “stiffer”, or more precisely, the relative actions of agonist and antagonist modulate the mechanical impedance (stiffness and damping) of the joint, as further discussed below.

A single muscle is considered to act as a brake when it operates on the lengthening region of the FV curve, but in a functional context such as in the current study, braking rarely occurs in isolation. Instead, it often follows motoring by the antagonist, in the current study resulting in a relatively high co-activation in the muscle pair during what might be traditionally called “braking” at the joint level. This is seen most prominently in the shoulder (Fig. 4C), but it is also visible in the elbow and wrist (Fig. 5B, 6B) although braking action of the elbow and wrist is complex (see below). As in the motor case, co-contraction can increase the impedance of the joint during braking action. This increased impedance is visible by observing the size of the joint-fv space which allows a visual reading of the instantaneous “stiffness” state of a joint. The examples above illustrate how the single-muscle view of “motor” or “brake” may be misleading at the joint level; one cannot strictly determine how much a muscle acts as a motor against the external load (e.g. limb inertia) versus against the antagonist. This ambiguity arises particularly in multibody systems where cross-joint interactions break the direct link between net muscle torque and joint movement (as is the case with the elbow; see below).

### The shape and direction of the joint-fv trajectory helps reveal the nature of underlying mechanical forces

To understand how joint-fv information links to underlying joint mechanics, we first note that the joint-fv trajectory is a description of muscle-produced torques, yet movement is governed by net joint torques. Consequently, the joint-fv trajectory may reveal situations where muscles produce a net torque, but the joint movement does not correspond to the direction of this net torque. While counter-intuitive, this reveals information about the underlying dynamics of the whole system, as explored further below.

For an isolated joint moving a constant purely inertial load, one expects a direct correspondence between acceleration and torque. However, in multibody systems, limb segments move not just when muscle forces are acting directly on them, but also when upstream (i.e. more proximal) segments are moving (Hollerbach & Flash, 1982). The angular velocities and accelerations of upstream joints give rise to local fictious forces (centrifugal, Coriolis, and Euler forces) which cause movement in downstream joints because joint angles are defined relative to the axis of the previous (rotating) segment. Additional multibody effects also arise when the changing joint positions causes the inertial properties of the limb to change.

To illustrate these principles, we focus on the forward reach as a representative example. For a hypothetical single-joint reach with no correction or homing-in, the joint follows a simple path through joint-FV space where increasing velocity (rightward motion in joint-FV space) corresponds to positive torque whereas leftward movement corresponds to negative torque (path a; Fig. 2D, 9B). During forward reaching, the joint-fv trajectory of the shoulder is a similar path, although it includes a “correction loop” (path b; Fig. 2D, 9B). This is expected, as the shoulder has no upstream joints and the change in the arm position is small enough not to cause major inertial changes. Net muscle torques at the elbow and wrist, however, do not correspond to their respective accelerations (Fig. 9E,F), instead diverging dramatically from paths a and b. Notably, at time ≈ 0.12s the elbow joint-fv trajectory veers sharply upwards/leftwards indicating increasing (positive) torque while decelerating (Fig. 9C; SI movie 3). The wrist shows a similar effect, though less pronounced (Fig. 6C; SI movie 3). This elbow joint-fv pattern is counterintuitive for two reasons: a) the joint is decelerating even when net muscle torque is positive and increasing (rather than accelerating as would be expected in a single-joint system), and b) activation of both muscles is decreasing while total elbow torque increases. Both of these observations suggest the presence of forces arising from outside the elbow muscles, specifically cross-joint interaction forces from the shoulder. These interaction forces can cause the total net torque on a joint to act in the opposite direction to the net muscle torque.

For the elbow, this cross-joint interaction occurs in three stages (Fig. 9C; SI movie 3): 1) At the onset, the elbow extensor torque drives positive (extension) elbow acceleration (rightward movement in quadrant I, shown by the thick green segment of the trajectory curve). This extension occurs despite opposing flexor torque caused by antagonism. 2) At time ≈ 0.09s (Fig. 9C point a) the flexor actions of the antagonist exceed the agonist action; the net torque action becomes flexion. However, despite the net flexor torque, the elbow continues to extend with increasing velocity until ∼0.11s (Fig 9C point b), and this continued extension is due to multibody forces which counteract and exceed the net muscle action.

Next, the joint begins to decelerate while the net muscle flexion torque increases, until ∼0.12s when the net flexion torque starts to decrease and the trajectory reverses direction back towards quadrant I (Fig. 9C point c-d). 3) From time ∼0.17s (Fig. 9C point e), the elbow crosses the x-axis again, returning to quadrant I, continuing to extend with increasing net extension torque, but counterintuitively, with diminishing speed. This behaviour indicates that while the net muscle torque would cause extension (extensor force exceeds flexor force) the net multibody torques work in the direction of flexion. During this stage, these multibody torques (mostly from shoulder deceleration) exceed elbow muscle torques, causing overall elbow deceleration. If not for these cross-joint effects, the elbow would follow a path similar to the shoulder (i.e. path b; Fig. 2D, 9B).

In light of the complex effects of cross-joint interactions described above, how useful are simple functional labels, such as motor and brake? We propose that these labels can be particularly useful in cases of clear separation between “sources and sinks” of mechanical energy (such as a muscle in a swimming fish transferring momentum to the water; e.g. Fish, 2010; Gerry & Ellerby, 2014) or a clear temporal separation between propulsion and braking (such as jumping versus landing in birds; e.g. Konow et al., 2012; Bishop et al., 2012). However, in other cases, the flow of mechanical energy between muscle and load may be less clear. In our reaching example above, the elbow extensor might be called a “motor” from the standpoint of mechanical energy; i.e. it shortens while producing force for the duration of its activity. This single-muscle perspective might suggest that its mechanical energy production contributes to the acceleration of a mass, but the muscle is mainly resisting combined forces of the antagonist and interaction torques from other joints. Hence, the “motor/brake” metaphors may not always be precise enough to capture the true mechanical state of muscles in the anatomical context of multiple joints and antagonistic muscle pairs.

### Co-contraction influences the shape of the joint-fv curve which may influence the response to perturbations, but further exploration is needed

Even when cross-joint effects do not dominate movement, the mechanical state of a joint can be influenced strongly by antagonist co-contraction. Given that co-contraction naturally occurs during reaching (e.g. Gribble et al., 2003) and may be an emergent response to excitation-contraction delays (Murtola & Richards, 2023), we aimed to investigate how co-contraction influences joint dynamics. Prior simulation work found that the intrinsic damping of a single muscle’s FV relationship helps to stabilise cyclic hopping motions (Haeufle et al., 2010). This intrinsic damping is most evident on the lengthening portion of the FV curve where an active muscle can absorb mechanical energy like a damper (e.g. from the recoil of a tendon following a landing impact; Konow et al., 2012). In theory, this damping ability becomes stronger during co-contraction because the antagonistic joint-fv curves add to steepen the local gradient, especially near zero velocity (Fig. 1B). This steepening indicates that any change in joint velocity is strongly counteracted by changes in the net muscle torque, which is most visible at the elbow for both normal reaching (Fig. 5B,C) and the perturbation (Fig. 9A). During the perturbation simulation, this enhanced damping is responsible for the rapid rebound which returns the joint torque nearly to pre-perturbation levels (Fig. 8B). We note that although the current work is not a rigorous perturbation study (see Van Wouwe et al., 2022), this simulated rapid rebound is a transient mechanical response which may augment other intrinsic stabilising features of muscle such as short-range stiffness (De Groote et al., 2017), and thus warrants further investigation beyond the current scope. Additionally, given that co-contraction has been observed as an adaptive response to novel force fields (Franklin et al., 2003) future work could address how co-contraction impacts task adaptation via changes in joint impedance arising from joint-FV dynamics.

### Despite the current limited scope, the joint-FV framework is not limited to simple “idealised” joints

For simplicity, our model (based on Murtola & Richards, 2023) only contains one agonist-antagonist pair per joint and neglects both tendons and bi-articular linkages. This level of complexity is sufficient for the current work because our main goal was to highlight conceptual insights gained by observing how FV properties interact with multibody dynamics. However, the conceptual joint-FV framework is not limited to simple models. As our framework utilises the mathematical dependency between muscle contraction speed and joint angular velocity to transform muscle-specific FV curves to the joint coordinate system, it can be generalised to account for additional curves from additional muscle actions. Furthermore, our framework allows muscles with unique properties (*vmax*, shape of the FV curve, moment arms, etc) to be summed (equivalently to Eq. 4) to create a multi-muscle, multi-articular joint-fv curve. Finally, the effect of tendons could be incorporated in future studies, as long as tendon properties are known. We note however that for the current work, tendons are unlikely to impact our results because the muscle forces (therefore tendon displacements) are relatively small. Furthermore, the current work only aims to introduce the framework and to highlight the rich information contained in joint-FV data. Hence, we feel that addition of further complexity (multiple muscles with varying properties) is beyond the current scope and will be addressed in subsequent studies.

Further to the above limitations, we did not investigate the effects of varying force-velocity properties. Importantly, the shape of the FV curve is known to vary significantly across species (e.g. in frogs; Astley, 2016) and among fibre types within a species (e.g. in mice; Kissane & Askew, 2024), and it has been found to strongly impact muscle function in musculoskeletal models (Charles et al., 2024). We note, however, that given the current focus on how generalised FV properties impact joint function, concerns about specific variation in FV parameters would not add meaningfully to the presented findings. However, a study of the influence of FV shape on joint-FV dynamics would be a fruitful area of future investigation, especially if joint-FV dynamics were to be analysed comparatively across species.

Finally, we acknowledge that Hill-type models lack stiffness and history-dependent effects (see Herzog, 2014; Nishikawa, 2020) and therefore have limited utility for studying transient behaviour such as general responses to perturbations. Hence, we restricted our perturbation to small forces (see methods) where the behaviour of Hill-type models remains reasonable. Moreover, we focused most of our analysis on movements where transient behaviour is not dominant. Regardless, more advanced models (e.g. Millard et al., 2024) would enable a thorough future study of how muscle impedance influences transient responses to perturbations. Although the joint-FV framework would need to be reworked for such models, the key concept still applies: multiple muscle-level states can be condensed into instantaneous joint-level representations by transforming them into the joint coordinate system.

### Summary and Conclusion

We developed a conceptual framework for visualising and describing how muscle force-velocity properties influence joint dynamics. In our investigation of simulated human reaching, we used joint-level force-velocity (joint-FV) plots to compare the dynamic function of the shoulder, elbow and wrist across three different reach trajectories. We found broad similarities across all joints and conditions, especially during the early phase of reaching. However, the elbow and wrist differed from the shoulder during the homing-in phase which showed a complex interplay of local muscle-driven torque and interaction torques due to shoulder movement. Additionally, we found that co-contraction creates a steepening of the joint-fv curve (compared to a single-muscle fv curve) which may confer increased damping and self-correction from small mechanical perturbations. More broadly, our new visualisation of joint-fv trajectories communicate joint kinetic function in the context of muscle activation and force-velocity dynamics that underlie this behaviour. We therefore propose that our joint-FV framework can be used to explain the intricate features seen in muscle data from musculoskeletal simulations enabling a better understanding of how intrinsic muscle properties limit the performance of dynamical systems.

## ACKNOWLEDGEMENTS

Firstly, we thank the late Professor Roger Woledge who shared his ideas with CR in 2013 at a curry house in S. Kensington where he speculated how co-contraction may steepen the fv curve at near isometric conditions. This was the core idea around which we’ve expanded and formalised our concept of a joint-level fv curve. We also thank Delyle Polet for insightful conversations around various aspects of this work during its preparation.

## DATA ACCESSIBILITY

All source code to run the simulations as well as simulation data, code for data analysis and code for figure generation will be provided in a github repository prior to publication.

## AUTHORS’ CONTRIBUTIONS

The original idea for the work was the joint conception of CR and TM. TM created the reaching model and modified it for current use. CR further modified the model to allow for perturbations. CR performed the numerical analyses, generated the figures and outlined the key findings. Both CR and TM contributed to writing the manuscript.

## SUPPLEMENTARY INFORMATION

SI Movie 1 Simulation of human reaching (control reaches). The animation shows the head, clavicle and right arm along with lines for muscle pathways. The grey sphere represents the hand and the blue sphere represents the reaching target. A series of three example reaches are shown: forward, sideways and cross-body. The total time for each reach is ∼0.5s. Note the MuJoCo simulation is rendered in 3D although the simulation is planar.

SI Movie 2 Simulation of human reaching (perturbed reaches). The animation is the same as SI Movie 1 except the simulations respond to mechanical perturbations for each of the three reaches (see text).

SI Movie 3 Joint FV space animation for the forward reach (no perturbation). The animation shows the temporal evolution of the joint-fv space for the shoulder (left), elbow (middle) and wrist (right). The inset on the shoulder joint-fv plot shows a stick figure animation of the right arm reaching for the target (red dot). The simulation time is shown with a time counter. The axes are identical as shown in Figs. 4, 5, 6.

SI Movie 4 Joint FV space animation for the sideways reach (no perturbation). See the description for SI Movie 3.

SI Movie 5 Joint FV space animation for the cross-body reach (no perturbation). See the description for SI Movie 3.

SI Movie 6 Joint FV space animation for the forward reach (with perturbation). See the description for SI Movie 3. Note the time counter turns red to indicate the period of the mechanical perturbation.

