## Supplementary material for "Visualising joint force-velocity properties in musculoskeletal models": Code&Data: mjcontrol.h

```
/***************************************************************
    MJCONTROL.H defines a structure of control variables 
    for SIMULATE_REACHES.CPP.

    Copyright 2022 Tiina Murtola/Royal Veterinary College

***************************************************************/


#pragma once

#include "stdio.h"

//---------------------------------- mjControl ---------------------------------------------

struct _mjControl
{

    /* FREELY ADJUSTIBLE VARIABLES */

    // ---- simulation parameters ----
    double sim_duration = 2.0;
    double error_tol = 0.001;//default 0.001

    // ---- target sequence control ----
    int ntargs = 4;  
    int next_targ = 0;
    double targ_dist = 0.2175;
    double reach_time = 1.0;        // value overwritten in code
    double reach_time_factor = 1; // value to be overwritten in main code

    // ---- control parameters ----

    // PD gains -- comment/uncomment or set manually as desired

    // Optima for different intrinsic muscle property setups
    // -- FMAX
    //double K[6] = { 5582.22, 1.0816 * sqrt(5582.22), 2050.08, 0.0051 * sqrt(2050.08),  7541.4, 0.0090 * sqrt(7541.4) };   
    //double K[6] = { 4342.35, 1.0816 * sqrt(4342.35), 5154.67, 0.0051 * sqrt(5154.67), 1622.29, 0.0090 * sqrt(1622.29) };

    // -- FMAX+O1
    //double K[6] = { 6920.38, 7.0322 * sqrt(6920.38), 328.13, 2.5314 * sqrt(328.13), 316.41, 0.3871 * sqrt(316.41) };
    //double K[6] = { 6919.67, 7.0008 * sqrt(6919.67), 325.00, 2.5313 * sqrt(325.00), 400.00, 0.3800 * sqrt(400.00) };

    // -- FVL+O1
    //double K[6] = { 7527.0, 6.3802 * sqrt(7527.0),  561.63, 2.3083 * sqrt(561.63), 223.45, 0.5001 * sqrt(223.45) };
    //double K[6] = { 7670.65, 6.2433 * sqrt(7670.65),  513.68, 2.498 * sqrt(513.68), 329.6, 0.4366 * sqrt(329.6) };

    // -- FMAX+O3				
    //double K[6] = { 38.33, 1.7957 * sqrt(38.33),  0.18, 0.5361 * sqrt(0.18), 2.96, 0.1417 * sqrt(2.96) };
    //double K[6] = { 67.89, 2.6026 * sqrt(67.89),  3.36, 0.9334 * sqrt(3.36), 15.03, 0.0895 * sqrt(15.03) };

    // -- FVL+O3
    //double K[6] = { 192.97, 1.3909 * sqrt(192.97), 671.47, 0.2971 * sqrt(671.47), 289.37, 0.126 * sqrt(289.37) };
    //double K[6] = { 853.76, 1.3384 * sqrt(853.76), 743.07, 0.4673 * sqrt(743.07), 270.68, 0.105 * sqrt(270.68) };//CHRIS USE THIS

    //double delay = 21;

    // -- computed torque (optima for passive stiffness scenarios)
    //double K[6] = { 1634.37, 1.6868 * sqrt(1634.37),  1598.33, 1.6840 * sqrt(1598.33),   1146.1, 2.0282 * sqrt(1146.1) };
    //double K[6] = { 77510.08, 6.7234 * sqrt(77510.08),  68802.36, 7.2061 * sqrt(68802.36),   57707.13, 7.8908 * sqrt(57707.13) };
    //double K[6] = { 77771.79, 6.7229 * sqrt(77771.79),  69121.53, 7.1996 * sqrt(69121.53),   57330.39, 7.8912 * sqrt(57330.39) };
    //double K[6] = { 71745.54, 6.6279 * sqrt(71745.54),  69600.11, 7.3660 * sqrt(69600.11),   62605.76, 8.3956 * sqrt(62605.76) };
    //double K[6] = { 75823.96, 6.3559 * sqrt(75823.96),  75872.38, 7.0780 * sqrt(75872.38),   76734.99, 7.6981 * sqrt(76734.99) };
    //double K[6] = { 86026.52, 8.5144 * sqrt(86026.52),  88755.10, 9.8127 * sqrt(88755.10),   78296.96, 9.4953 * sqrt(78296.96) };
    //double K[6] = { 76312.34, 6.4059 * sqrt(76312.34),  79309.97, 7.4841 * sqrt(79309.97),   73724.74, 6.4667 * sqrt(73724.74) };//default
    //double K[6] = { 29628.14, 8.6073 * sqrt(29628.14),  37985.79, 8.3941 * sqrt(37985.79),   49588.35, 13.2613 * sqrt(49588.35) };
    //double K[6] = { 66093.67, 7.5673 * sqrt(66093.67),  74609.96, 6.8518 * sqrt(74609.96),   94330.22, 5.4441 * sqrt(94330.22) };


    //------------------SLOW REACH 3RD ORDER ACT DYNAMICS + FVL + computed torque------------------------
    //parameters re-optimised by Tiina, 18.12.23
    //optimised for reach factor = 1.712 (control)
    // int act_mode = 0;//0 for 3rd order, 1 for 1st order
    // double delay = 28;
    // double K[6] = { 1634.37, 1.6868 * sqrt(1634.37), 1598.33, 1.684 * sqrt(1598.33), 1146.1, 2.0282 * sqrt(1146.1) };

    //------------------FAST REACH 3RD ORDER ACT DYNAMICS + FVL + computed torque------------------------
    //parameters re-optimised by Tiina, 18.12.23
    //optimised for reach factor = 0.43 (4x speed)
    int act_mode = 0;//0 for 3rd order, 1 for 1st order
    double delay = 25;
    double K[6] = { 684.45, 1.6064 * sqrt(684.45), 3685.82, 1.5886 * sqrt(3685.82), 6423.55, 1.1522 * sqrt(6423.55) }; 

    //------------------SLOW REACH 1ST ORDER ACT DYNAMICS + FVL + computed torque------------------------
    //parameters re-optimised by Tiina, 18.12.23
    //optimised for reach factor = 1.712 (control)
    // int act_mode = 1;//0 for 3rd order, 1 for 1st order
    // double delay = 6;
    // double K[6] = { 95236.09, 2.0853 * sqrt(95236.09), 64175.31, 2.3015 * sqrt(64175.31), 19537.03, 5.19 * sqrt(19537.03) };                    

    //------------------FAST REACH 1ST ORDER ACT DYNAMICS + FVL + computed torque------------------------ 
    //parameters re-optimised by Tiina, 18.12.23
    //optimised for reach factor = 0.43 (4x speed)
    // int act_mode = 1;//0 for 3rd order, 1 for 1st order
    // double delay = 6;
    // double K[6] = { 34955.02, 4.5685 * sqrt(34955.02), 47609.59, 5.0843 * sqrt(47609.59), 85439.72, 3.9995 * sqrt(85439.72) };    

    /* ADJUSTIBLE WITHIN XML GEOMTRY LIMITS */

    double initPose[3] = { 20, 80, 20 };
    double shoulder_pos[3] = { 0.0, 0.0, 0.0 };  // values overwritten in code


   
    /* KEY INITIALISATIONS - DO NOT TOUCH */

    // --------- force variables ------------
    mjtNum* gain_len;
    mjtNum* gain_vel;

    // loop control variables
    bool use_predicted_data = 0;
    bool update_control = 1;
    double error_norm = 1.0;
    mjtNum cum_error_res = 0.0;
    mjtNum cum_error_d = 0.0;

    // task variables
    bool success = 0;
    double time_succ = -1.0;

    // target vectors
    mjtNum targ_xpos[3];            // target position in Cartesian coordinates
    mjtNum targ_xvel[3];             // target velocity in Cartesian coordinates
    mjtNum* targ_qvel;   
    mjtNum* targ_qacc;

    // error vectors
    mjtNum x_error[3];               // position error (wrt target) in cartesian position
    mjtNum x_error_init[3];          // position error of initial pose
    mjtNum x_error_d[3];             // position error wrt desired position
    mjtNum* q_error;                 // position error in joint coordinates
    mjtNum* qvel_error;              // velocity error in joint coordinates


    // prediction
    mjtNum* predicted_qvel;
    mjtNum predicted_xpos[3];
    mjtNum x_error_pred[3];

    // Jacobians & related matrices
    mjtNum* Jcb;                   // end-effector Jacobian
    mjtNum* Jpl;                    // pseudo-inverse of jacp

    // torques & forces
    mjtNum* PDmagn;  
    mjtNum* computed_torque;
    mjtNum* PDpert;//addedf by CTR for injecting a perturbation to the PD control vector
    mjtNum* PDprev;//PDmagn at previous time step
    mjtNum* computed_torqueFF;//computed torque opened from saved file from a Feedback simulation


    void init(const int nv)
    {
        gain_len = new mjtNum[2 * nv];  // needs fixing if number of muslces is not 2x dofs
        gain_vel = new mjtNum[2 * nv];

        targ_qvel = new mjtNum[nv];     
        targ_qacc = new mjtNum[nv];

        q_error = new mjtNum[nv];
        qvel_error = new mjtNum[nv];

        predicted_qvel = new mjtNum[nv];

        Jcb = new mjtNum[3 * nv];
        Jpl = new mjtNum[nv * 3];
       
        PDmagn = new mjtNum[nv];
        computed_torque = new mjtNum[nv];
        PDpert = new mjtNum[nv];
        PDprev = new mjtNum[nv];
        computed_torqueFF = new mjtNum[nv];
    }

    void del()
    {
        delete[] gain_len;
        delete[] gain_vel;

        delete[] targ_qvel;
        delete[] targ_qacc;

        delete[] q_error;
        delete[] qvel_error;

        delete[] predicted_qvel;

        delete[] Jcb;
        delete[] Jpl;

        delete[] PDmagn;
        delete[] computed_torque;
        delete[] PDpert;
        delete[] PDprev;
        delete[] computed_torqueFF;
    }
};
typedef struct _mjControl mjControl;
```
