## Supplementary material for "Visualising joint force-velocity properties in musculoskeletal models": Code&Data: reachingToolbox.h

```
/***************************************************************
    REACHINGTOOLBOX.H defines auxiliary functions and custom 
    callbacks for SIMULATE_REACHES.CPP.

    Copyright 2023 Tiina Murtola/Royal Veterinary College

***************************************************************/

#pragma once
#include <cmath>
#include <math.h>
//#include "LinAlg.h"
#include "linalg.h"
//#include "alglibinternal.h"
//#include "alglibmisc.h"
//#include "ap.h"
#include <algorithm>
// file input/output
#include <iostream>
#include <iomanip>
#include <fstream>
#include <string>
//#include <sstream>
#include <vector>

extern mjControl c;
extern mjData* dpred;


/* MUSCLE FUNCTIONS */

double FLgain(double l, double a, double b, double c)
{
    double Fgain;
    Fgain = exp(-pow(abs((pow(l, b) - 1) / c), a));

    return Fgain;
}

double FVgain(double v, double a, double b, double c, double vmax)
{
    double vnorm = -v / vmax;    //minus sign due to MuJoCo's sign convention
    double Fgain;

    if (vnorm > 0)
        Fgain = (1 - vnorm) / (1 + a * vnorm);
    else
        Fgain = b - (b - 1) * (1 + vnorm) / (1 - a * c * vnorm);

    return Fgain;
}

//returns the size of a csv text file in rows and columns
int sizeofCSV(int* rows, int* cols, FILE* file){
    
    int ch, r, ic;
    
    ch = r = 0; ic = 1;
    
    while(!feof(file))
    {
       ch = fgetc(file);
        if((r == 0) && (ch == ','))
            ic++;
        if(ch == '\n')
            r++;
    }
    
    //very important - return file pointer to start of file
    rewind(file);
    
    //write output
    *rows = r; *cols = ic;
    return r;
}


void optimalLengthFromCurrent(mjModel* m, const mjData* d)
{
    /* Saves current length of actuator in m->actuator_user
    * for use as optimal muscle length in muscleBias and
    * muscleGain functions. Also intialises velocity and
    * length gains for each muscle in mjcontrol struct c.
    */
    for (int actID = 0; actID < m->nu; actID++)
    {
        m->actuator_user[actID] = d->actuator_length[actID];

        // initialise muscle length and velocity gains for potential printing
        c.gain_len[actID] = FLgain((d->actuator_length[actID]) / m->actuator_user[actID], m->actuator_gainprm[actID * 10 + 1],
            m->actuator_gainprm[actID * 10 + 2], m->actuator_gainprm[actID * 10 + 3]);

        c.gain_vel[actID] = FVgain(d->actuator_velocity[actID] / m->actuator_user[actID], m->actuator_gainprm[actID * 10 + 4],
            m->actuator_gainprm[actID * 10 + 5], m->actuator_gainprm[actID * 10 + 6],
            m->actuator_gainprm[actID * 10 + 7]);
    }
}


mjtNum muscleActivation1stOrder(const mjModel* m, const mjData* d, int id)
{
    /* Computes and returns act_dot (for d->act_dot) for first order nonlinear
        activation dynamics. Intended for use as activation callback
        in simulation pipeline. To use, set
            mjcb_act_dyn = muscleActivation1stOrderNL;
       and check m->actuator_dynprm set in model file.
     */

    mjtNum act_dot;

    double tau;
    // time constant
    if (d->ctrl[id] > d->act[id])
        tau = m->actuator_dynprm[id * 10] * (0.5 + 1.5 * d->act[id]);
    else
        tau = m->actuator_dynprm[id * 10 + 1] / (0.5 + 1.5 * d->act[id]);

    // first-order activation dynamics
    act_dot = (d->ctrl[id] - d->act[id]) / tau;

    return act_dot;
}

mjtNum muscleActivation3rdOrder(const mjModel* m, const mjData* d, int id)
{
    /* Computes and returns act_dot (for d->act_dot) for third order
    *  acivation dynamics. Intended to be use as activation callback 
    *  in simulation pipeline. To use, set
    *    mjcb_act_dyn = muscleActivation3rdOrder;
    *  and check m->actuator_dynprm set in model file.
    */
    mjtNum act_dot;

    d->userdata[3 * id] = d->userdata[3 * id] + m->opt.timestep / m->actuator_dynprm[id * 10 + 4] * d->ctrl[id]
        - m->opt.timestep / m->actuator_dynprm[id * 10 + 4] * (m->actuator_dynprm[id * 10 + 5] 
            + (1 - m->actuator_dynprm[id * 10 + 5]) * d->ctrl[id]) * d->userdata[3 * id];

    d->userdata[3 * id + 1] = d->userdata[3 * id + 1] + m->opt.timestep / m->actuator_dynprm[id * 10 + 2] * d->userdata[3 * id]
        - m->opt.timestep / m->actuator_dynprm[id * 10 + 2] * (m->actuator_dynprm[id * 10 + 3] 
            + (1 - m->actuator_dynprm[id * 10 + 3]) * d->userdata[3 * id]) * d->userdata[3 * id + 1];

    d->userdata[3 * id + 2] = d->userdata[3 * id + 2] + m->opt.timestep / m->actuator_dynprm[id * 10] * d->userdata[3 * id + 1]
        - m->opt.timestep / m->actuator_dynprm[id * 10] * (m->actuator_dynprm[id * 10 + 1] 
            + (1 - m->actuator_dynprm[id * 10 + 1]) * d->userdata[3 * id + 1]) * d->userdata[3 * id + 2];

    act_dot = d->userdata[3 * id + 1] / m->actuator_dynprm[id * 10] - (m->actuator_dynprm[id * 10 + 1] + (1 - m->actuator_dynprm[id * 10 + 1]) * d->userdata[3 * id + 1]) * d->act[id] / m->actuator_dynprm[id * 10];


    return act_dot;
}


mjtNum muscleGainFLV(const mjModel* m, const mjData* d, int id)
{
    /* Returns muscle force from contractile element excluding the effects to dynamic activation.
       Intended to use as muscle gain callback in simulation pipeline. To use, set
            mjcb_act_gain = muscleGainFLV;
       and check m->actuator_gainprm.
    */

    mjtNum ForceGain;

    // force gain from F-L curve
    c.gain_len[id] = FLgain((d->actuator_length[id]) / m->actuator_user[id], m->actuator_gainprm[id * 10 + 1],
        m->actuator_gainprm[id * 10 + 2], m->actuator_gainprm[id * 10 + 3]);

    // force gain from F-V curve
    c.gain_vel[id] = FVgain(d->actuator_velocity[id] / m->actuator_user[id], m->actuator_gainprm[id * 10 + 4],
        m->actuator_gainprm[id * 10 + 5], m->actuator_gainprm[id * 10 + 6],
        m->actuator_gainprm[id * 10 + 7]);   // velocity in the same units as s.vmax

    // total gain = Fmax*FVgain*FLgain (activation gain in a separate function)
    ForceGain = m->actuator_gainprm[id * 10] * c.gain_vel[id] * c.gain_len[id]; // default assumption of 10 gain parameters per actuator

    return ForceGain;
}


mjtNum muscleGainConst(const mjModel* m, const mjData* d, int id)
{
    /* Returns muscle force from contractile element excluding the effects to dynamic activation.
       Intended to use as muscle gain callback in simulation pipeline. To use, set
            mjcb_act_gain = muscleGainConst;
       and check m->actuator_gainprm.
    */

    mjtNum ForceGain;

    ForceGain = m->actuator_gainprm[id * 10];

    return ForceGain;
}

mjtNum muscleBias(const mjModel* m, const mjData* d, int id)
{
    /* Returns passive force from parallel elastic element.
       Intended to use as a muscle bias callback in simulation pipeline. To use, set
            mjcb_act_bias = muscleBias;
       and check m->actuator_biasprm.
    */

    mjtNum ForceBias;
    double delta_l_rel;

    delta_l_rel = d->actuator_length[id] / m->actuator_user[id] - m->actuator_biasprm[id * 10 + 2];

    if (delta_l_rel > 0)
        ForceBias = m->actuator_biasprm[id * 10] * (exp(m->actuator_biasprm[id * 10 + 1] * delta_l_rel) - 1);
    else
        ForceBias = 0.0;

    return ForceBias;
}


/* CONTROL FUNCTIONS */

void psinv(mjtNum* res, mjtNum* J, const int n)
{
    /* pseudo-inverse of J through SVD */

    alglib::real_2d_array a, u, vt, sinv, temp, ainv;
    alglib::real_1d_array w;

    int m = 3;
    int min_dim = std::min(m, n);

    //// Initialize arrays
    a.setlength(m, n);
    u.setlength(m, m);
    vt.setlength(n, n);
    w.setlength(min_dim);
    sinv.setlength(n, m);
    temp.setlength(n, m);
    ainv.setlength(n, m);

    for (int i = 0; i < m; i++)
    {
        for (int j = 0; j < n; j++)
        {
            a[i][j] = J[i * n + j];
            sinv[j][i] = 0.0;
        }
    }

    alglib::setglobalthreading(alglib::parallel);
    alglib::rmatrixsvd(a, m, n, 2, 2, 2, w, u, vt);

    for (int i = 0; i < min_dim; i++)
    {
        if (abs(w[i]) > 0.00001)
            sinv[i][i] = 1 / w[i];
        else
            sinv[i][i] = 0;
    }

    alglib::rmatrixgemm(n, m, n, 1.0, vt, 0, 0, 1, sinv, 0, 0, 0, 0.0, temp, 0, 0);
    alglib::rmatrixgemm(n, m, m, 1.0, temp, 0, 0, 0, u, 0, 0, 1, 0.0, ainv, 0, 0);

    for (int i = 0; i < n; i++)
    {
        for (int j = 0; j < m; j++)
            res[i * m + j] = ainv[i][j];
    }
}


void computePDmagn(const mjModel* m, mjData* d)
{
    int grpid = mj_name2id(m, mjOBJ_SITE, "grip");
    int trgid = mj_name2id(m, mjOBJ_SITE, "target");

    // -- position error in cartesian coordinates wrt target
    c.x_error[0] = m->site_pos[3 * trgid] - d->site_xpos[3 * grpid];
    c.x_error[1] = m->site_pos[3 * trgid + 1] - d->site_xpos[3 * grpid + 1];
    c.x_error[2] = 0;  // 2D motion    

    if (c.use_predicted_data)
    {
        c.x_error_pred[0] = m->site_pos[3 * trgid] - c.predicted_xpos[0];
        c.x_error_pred[1] = m->site_pos[3 * trgid + 1] - c.predicted_xpos[1];
        c.x_error_pred[2] = 0;
        // -- compute the Jacobian
        mj_jacSite(m, dpred, c.Jcb, NULL, grpid);
    }
    else
        // -- compute the Jacobian
        mj_jacSite(m, d, c.Jcb, NULL, grpid);

    // -- pseudo-inverse of the Jacobian
    psinv(c.Jpl, c.Jcb, m->nv);

    // -- compute positional and velocity errors
    mjtNum vk;
    mjtNum wk;

    if (c.use_predicted_data)
    {
        double pred_time = d->time + m->opt.timestep * (int)c.delay;
        double tau = pred_time / c.reach_time;
        if (pred_time > c.reach_time)
        {
            vk = 0.0; wk = 1.0;
        }
        else
        {
            vk = 1 / c.reach_time * abs(60 * pow(tau, 3) - 30 * pow(tau, 4) - 30 * pow(tau, 2));
            wk = abs(15 * pow(tau, 4) - 6 * pow(tau, 5) - 10 * pow(tau, 3));
        }
        mju_addScl3(c.x_error_d, c.x_error_pred, c.x_error_init, (wk - 1));
    }
    else
    {
        double tau = d->time / c.reach_time;
        if (d->time > c.reach_time)
        {
            vk = 0.0; wk = 1.0;
        }
        else
        {
            vk = 1 / c.reach_time * abs(60 * pow(tau, 3) - 30 * pow(tau, 4) - 30 * pow(tau, 2));
            wk = abs(15 * pow(tau, 4) - 6 * pow(tau, 5) - 10 * pow(tau, 3));
        }
        mju_addScl3(c.x_error_d, c.x_error, c.x_error_init, (wk - 1.0));
    }

    mju_mulMatVec(c.q_error, c.Jpl, c.x_error_d, m->nv, 3);

    mju_scl3(c.targ_xvel, c.x_error_init, vk);
    mju_mulMatVec(c.targ_qvel, c.Jpl, c.targ_xvel, m->nv, 3);


    // -- velocity error in joint coordinates
    if (c.use_predicted_data)
        mju_sub(c.qvel_error, c.targ_qvel, c.predicted_qvel, m->nv);
    else
        mju_sub(c.qvel_error, c.targ_qvel, d->qvel, m->nv);

    for (int qID = 0; qID < m->nv; qID++)
    {
        c.PDmagn[qID] = c.K[2 * qID] * c.q_error[qID] + c.K[2 * qID + 1] * c.qvel_error[qID];//modified by cTR to add control perturbation PDpert
    }
}

void muscleControlTargetPD(const mjModel* m, mjData* d)
{
    /* Computes input control signal for muscles based on the error between target and grip site
    positions using basic PD control approach.
    */

    if (!c.update_control)
    {
        return;
    }

    computePDmagn(m, d);

    double act_torque;
    double max_torque;
    int best_actuator;
    double max_antg_torque;
    int antg_actuator;
    double s = 1.0;

    mju_zero(d->ctrl, m->nu);

    for (int qID = 0; qID < m->nv; qID++)
    {
        max_torque = 0.0;
        max_antg_torque = 0.0;
        for (int actID = 0; actID < m->nu; actID++)
        {
            if (c.use_predicted_data)
                act_torque = d->actuator_moment[actID * m->nv + qID] * muscleGainFLV(m, dpred, actID);
            else
                act_torque = d->actuator_moment[actID * m->nv + qID] * muscleGainFLV(m, d, actID);

            if (act_torque * c.PDmagn[qID] > 0.0 && abs(act_torque) > abs(max_torque)) // highest torque in the right direction
            {
                best_actuator = actID;
                max_torque = act_torque;
            }
            else if (act_torque * c.PDmagn[qID] < 0.0 && abs(act_torque) > abs(max_antg_torque)) // highest torque in the opposite direction
            {
                antg_actuator = actID;
                max_antg_torque = act_torque;
            }
        }

        if (abs(max_torque) > 0)
        {
            d->ctrl[best_actuator] = abs(c.PDmagn[qID] / max_torque);
        }
    }
}

void muscleControlTargetCompTorq(const mjModel* m, mjData* d)
{
    /*Computes input control signal for muscles based on the error between target and grip site
    positions using the computed torque approach.
    */

    if (!c.update_control)
    {
        return;
    }

    computePDmagn(m, d);
    mjtNum PDmagAugmented[m->nv];//this will be the sum c.PDmagn + c.PDpert
    mju_add(PDmagAugmented, c.PDmagn, c.PDpert, m->nv);

    if (c.use_predicted_data)
        //mj_mulM(m, dpred, c.computed_torque, c.PDmagn);
        mj_mulM(m, dpred, c.computed_torque, PDmagAugmented);
    else
        //mj_mulM(m, d, c.computed_torque, c.PDmagn);
        mj_mulM(m, d, c.computed_torque, PDmagAugmented);//incorporate control signal perturbation into computed torque
    mju_addTo(c.computed_torque, d->qfrc_bias, m->nv);


    double act_torque;
    double max_torque;
    int best_actuator;

    mju_zero(d->ctrl, m->nu);

    //printf("torques: ");
    for (int qID = 0; qID < m->nv; qID++)
    {
        max_torque = 0.0;
        for (int actID = 0; actID < m->nu; actID++)
        {
            if (c.use_predicted_data)
                act_torque = d->actuator_moment[actID * m->nv + qID] * muscleGainFLV(m, dpred, actID);
            else
                act_torque = d->actuator_moment[actID * m->nv + qID] * muscleGainFLV(m, d, actID);

            if (act_torque * c.computed_torque[qID] > 0.0 && abs(act_torque) > abs(max_torque)) // highest torque in right direction
            {
                best_actuator = actID;
                max_torque = act_torque;
            }
        }

        if (abs(max_torque) > 0)
        {
            d->ctrl[best_actuator] = abs(c.computed_torque[qID] / max_torque);
            if (d->ctrl[best_actuator] > 1.0)
                d->ctrl[best_actuator] = 1.0;

        }

    }

}

//obsolete function used to open excitation data and save into control vector u
//use ffComputedTorqueFromFile instead
void ffControlFromFile(const mjModel* m, mjtNum* u, FILE* file) 
{
    //find size of ctrl file
    int rows;
    int cols;
    sizeofCSV(&rows, &cols, file);
    int r,ic,ret; //ret value can be used for error checking
    
    if (cols > m->nu)
        cols = m->nu;
    //printf("rows and columns %i, %i\n", rows, cols); 
    //fill the arrays
    for(r=0;r<rows;r++)
    {
        //get each value followed by a comma
        for(ic=0;ic<cols-1;ic++)
        {
            ret = fscanf(file,"%lf,",&u[r*m->nu + ic]);
        }
        
        if(r < rows-1)  //get last value followed by newline
            ret = fscanf(file,"%lf\n",&u[r*m->nu + ic]);
        else            //get last value at end of file
            ret = fscanf(file,"%lf",  &u[r*m->nu + ic]);
        
    }
    
    //close the file now that we are finished
    fclose(file);
}

void ffComputedTorqueFromFile(const mjModel* m, mjtNum* ctrq, FILE* file) 
{
    //find size of ctrl file
    int rows;
    int cols;
    sizeofCSV(&rows, &cols, file);
    int r,ic,ret; //ret value can be used for error checking
    
    if (cols > m->nv)
        cols = m->nv;
    //printf("rows and columns %i, %i\n", rows, cols); 
    //fill the arrays
    for(r=0;r<rows;r++)
    {
        //get each value followed by a comma
        for(ic=0;ic<cols-1;ic++)
        {
            ret = fscanf(file,"%lf,",&ctrq[r*m->nv + ic]);
        }
        
        if(r < rows-1)  //get last value followed by newline
            ret = fscanf(file,"%lf\n",&ctrq[r*m->nv + ic]);
        else            //get last value at end of file
            ret = fscanf(file,"%lf",  &ctrq[r*m->nv + ic]);
        
    }
    
    //close the file now that we are finished
    fclose(file);
}

//do Feedforward (FF) control based on the control data u loaded from .csv file which comes from
//ctrl (excitation) data from a successful feedback control trial saved previously
//here is where internal perturbations are processed via c.PDpert --> acceleration perturbation
//mapped into muscle controls.
//r is the simulation timestep counter

void muscleControlFF(const mjModel* m, mjData* d)
{
    //return;
    double act_torque;
    double max_torque;
    int best_actuator;
    mjtNum ctrlFF;

    //augmented c.computed torque which has the current (unperturbed) torque added with c.PDpert
    //create and use c.computed_torque_FF i.e. c.computed toruqe from NO perturbation, but at the current timestep
    //easiest way is save c.computed_torque to file then --- this can be used as FF command instead of excitations!
    //THEN remove line 520/21 and 481 in main code ///ALSO- currently PDpert in FB mode works identically, but to make code easier to read, put addition of c.PDpert innside the muscleControlTargetCompTorq, i.e in line  409
    mj_mulM(m, d, c.computed_torque, c.PDpert);//Multiply mass matrix by PD vector which are accelerations
    //mjtNum computed_torque_Augmented[m->nv];
    //mju_add(computed_torque_Augmented, c.computed_torque, c.computed_torqueFF, m->nv);//compute torque based on sum of unperturbed FF signal and perturbation control signal
    //mju_copy(c.computed_torque, computed_torque_Augmented, m->nv);//overwrite c.computed_torque
    //--- Map computed torque into muscle excitations
    for (int qID = 0; qID < m->nv; qID++)
    {
        max_torque = 0.0;
        for (int actID = 0; actID < m->nu; actID++)
        {
            act_torque = d->actuator_moment[actID * m->nv + qID] * muscleGainFLV(m, d, actID);

            if (act_torque * c.computed_torque[qID] > 0.0 && abs(act_torque) > abs(max_torque)) // highest torque in right direction - note moment arm is signed
            {
                best_actuator = actID;
                max_torque = act_torque;
            }
        }

        if (abs(max_torque) > 0)
        {
            ctrlFF = d->ctrl[best_actuator];//FF control value for given muscle prior to perturbation
            d->ctrl[best_actuator] = abs(c.computed_torque[qID] / max_torque) + ctrlFF;//scales computed torque with max torque so it's 0-1
            if (d->ctrl[best_actuator] > 1.0)
                d->ctrl[best_actuator] = 1.0;

        }

    }
}


/* TASK CONTROL FUNCTIONS */

void setInitialPose(const mjModel* m, mjData* d, double initPose[])
{
    for (int qID = 0; qID < m->nq; qID++)
        d->qpos[qID] = initPose[qID] / 180 * mjPI;
    mj_forward(m, d);
}

void importTargets(std::ifstream& targetfile, std::vector<std::pair<double, double>>& targ_list)
{
    std::string line;
    std::string::size_type sz;
    std::string::size_type st;

    int numberOfLines = 0;
    int varInd = 0;
    double x_targ[2];
    while (std::getline(targetfile, line))
    {
        st = 0;
        for (int varInd = 0; varInd < 2; varInd++)
        {
            x_targ[varInd] = std::stof(line.substr(st), &sz);
            sz++; // correction due to separator ";\t"
            st += sz;
        }
        targ_list.push_back(std::make_pair(x_targ[0], x_targ[1]));
        ++numberOfLines;
    }

    std::cout << "Reading targets completed with " << numberOfLines << " lines of data.\n";
    c.ntargs = numberOfLines;
}

//from CTR, 11.23 - allows user to append misc parameter values separated by commas after each target pair as follows:
// xcoord;ycoord;<value1,value2,value3>
void importMiscFromTargetFile(std::ifstream& targetfile, mjtNum* misc_parameters, int target_number, int n_parameters)
{

    std::string line;
    std::string strBtwdelimiters;
    std::string::size_type sz;
    std::string::size_type st;

    int iline = 0;
    while (std::getline(targetfile, line))
    {
        //extract text between <> delimiters
        unsigned first = line.find("<");
        unsigned last = line.find_last_of(">");
        strBtwdelimiters = line.substr(first + 1, last-first - 1);

        if (iline == target_number)
        {
            st = 0;
            for (int iparam = 0; iparam < n_parameters; iparam++)
            {
                misc_parameters[iparam] = std::stof(strBtwdelimiters.substr(st), &sz);
                sz++; // correction due to separator ";\t"
                st += sz;
                //printf("test vector %f\n", misc_parameters[iparam]);
            }

        }

        ++iline;
    }
}


bool nextTarget(mjModel* m, const mjData* d, const double targ_xpos[2])
{
    int grpid = mj_name2id(m, mjOBJ_SITE, "grip");
    int trgid = mj_name2id(m, mjOBJ_SITE, "target");
    bool next_target_exists = 0;

    if (c.next_targ < c.ntargs)
    {
        c.targ_xpos[0] = targ_xpos[0];
        c.targ_xpos[1] = targ_xpos[1];
        c.targ_xpos[2] = 0.0;

        c.x_error_init[0] = c.targ_xpos[0] - d->site_xpos[3 * grpid];
        c.x_error_init[1] = c.targ_xpos[1] - d->site_xpos[3 * grpid + 1];
        c.x_error_init[2] = 0.0;

        double targ_dist = sqrt(c.x_error_init[0] * c.x_error_init[0] + c.x_error_init[1] * c.x_error_init[1]);
        c.reach_time = abs (c.reach_time_factor) * pow(targ_dist, 1.0 / 3.0);//reach_time_factor is from misc_parameters loaded from target file
        //c.reach_time = 1.712 * pow(targ_dist, 1.0 / 3.0);//normal speed reach
        //c.reach_time = 0.57 * pow(targ_dist, 1.0 / 3.0);//scaled speed to be 1/3 normal speed - it can be as low as 0.45

        next_target_exists = 1;

        m->site_pos[3 * trgid] = c.targ_xpos[0];
        m->site_pos[3 * trgid + 1] = c.targ_xpos[1];
        printf("\ntarget distance %f, reach time %f\n", targ_dist, c.reach_time);
    }
    else
        next_target_exists = 0;

    return next_target_exists;
}


void updateErrors(const mjModel* m, mjData* d)
{
    c.error_norm = mju_norm3(c.x_error);

    if (c.error_norm < c.error_tol)
    {
        c.success = 1;
        if (c.time_succ < 0) // first contact
        {
            c.time_succ = d->time;
            c.cum_error_res = 0.0;
        }
    }

    c.cum_error_res += c.error_norm;

    double tau;
    double wk;
    mjtNum temp_error_d[3];

    if (!c.use_predicted_data)  // x_error_d corresponds to current timestep
        c.cum_error_d += mju_norm3(c.x_error_d);
    else if (d->time > c.reach_time) // wk=1, so error from desired = error from target
        c.cum_error_d += c.error_norm;
    else
    {
        tau = (d->time - m->opt.timestep) / c.reach_time; //time at time of xd computation
        wk = abs(15 * pow(tau, 4) - 6 * pow(tau, 5) - 10 * pow(tau, 3));

        mju_addScl3(temp_error_d, c.x_error, c.x_error_init, (wk - 1));
        c.cum_error_d += mju_norm3(temp_error_d);
    }
}

//function added by CTR, 11.23 to produce various modes of perturbation depending on misc user parameters passed through 
//the target file.  It can be modified later for force field perturbations if needed
void perturbationGen(const mjModel* m, mjData* d, mjtNum* misc_parameters, int current_timestep)
{
    int grpid = mj_name2id(m, mjOBJ_SITE, "grip");
    mjtNum xhand[3];//hand position in cartesian, just for checking
    mjtNum xhand_prev[3];
    mjtNum vhand_unit[3];//cartesian hand velocity, just for checking
    for (int i = 0; i < 3; i++) 
    {
        xhand[i] = d->site_xpos[3 * grpid + i];
    }

    int body_pert = 3;//-- for external force perturbation to hand
    int pert_start = round(c.reach_time * misc_parameters[4] / (m->opt.timestep) );//start time of perturbation (in samples)
    int pert_end = pert_start + misc_parameters[5];
    mjtNum pert_magn = misc_parameters[2];
    //-- for external/internal perturbation 
    //external: find the instantaneous vector for cartesian hand velocity
    //internal: calculate an instantaneous vector which is the instantanous rate of change of PD control
    //since this is 3 joints, it's a 3D vector
    //perturbation will be a 3D vector added to velocity or control vector at 4 different angles (N,S,E,W) with respect to main vector
    mjtNum vec_pert[3];//unit vector with respect to hand velocity (external) or pd velocity (internal)
    mjtNum vec_pert_scaled[3];//to be scaled with the magnitude of the perturbation
    mjtNum pdpert_magn = misc_parameters[2];//magnitude of the PD control perturbation
    mjtNum pert_quat[4];//quaternion which will rotate the perturbation vector to the appropriate orientation

    if ( (current_timestep >= pert_start) && (current_timestep < pert_end) ) 
    { 
        if (misc_parameters[1] == 0) //external perturbation = step force applied at an angle with respect to current hand cartesian velocty
        {
            //find cartesian velocity of hand, normalise it then rotate it to get aixs of perturbation
            mjtNum xvel_unit[3] = {1, 0, 0};
            //mju_copy3(xvel_unit, c.targ_xvel);//hand velocity towards target in cartesian coordinates
            mju_copy3(xvel_unit, c.x_error);//hand velocity towards target in cartesian coordinates - should be in direction of error
            mju_normalize(xvel_unit, 3);
            mju_scl3(vec_pert_scaled, xvel_unit, pert_magn);
            mjtNum pert_axis[3] = {0, 0, 1};// will rotate PD control vector about Z axis - equivalent to perturbing shoulder and
            //elbow control signals only because control happens mostly in the shoulder-elbow plane
            //printf("PD velocity %f, %f, %f \n", pdvel[0], pdvel[1], pdvel[2]);
            mjtNum pert_angle = misc_parameters[3];//angle to rotate the perturbation vector wrt current PD control vector orientation
            mju_axisAngle2Quat(pert_quat, pert_axis, pert_angle);
            mju_rotVecQuat(vec_pert, vec_pert_scaled, pert_quat);
            printf("External perturbation of %f N delivered at time =  %f\n", pert_magn, d->time);
            for (int i = 0; i < 3; i ++) //first 3 values are XYZ components of xfrc_applied
            {
                d->xfrc_applied[body_pert * 6 + i] = vec_pert[i];//vec_pert[i];
            }
            //printf("vec pert %f, %f, %f\n", xvel_unit[0], xvel_unit[1], xvel_unit[2]);
            //printf("hand pos %f, %f, %f\n", xhand[0], xhand[1], xhand[2]);
        }
        else if (misc_parameters[1] == 1) //internal perturbation
        {
            //compute "PD velocity"
            mjtNum pdvel[3];//rate of change of pd vector
            mju_sub(pdvel, c.PDmagn, c.PDprev, 3);
            mju_normalize(pdvel, 3);
            mju_scl3(vec_pert_scaled, pdvel, pdpert_magn);
            mjtNum pert_axis[3] = {0, 0, 1};// will rotate PD control vector about Z axis - equivalent to perturbing shoulder and
            //elbow control signals only because control happens mostly in the shoulder-elbow plane
            //printf("PD velocity %f, %f, %f \n", pdvel[0], pdvel[1], pdvel[2]);
            mjtNum pert_angle = misc_parameters[3];//angle to rotate the perturbation vector wrt current PD control vector orientation
            mju_axisAngle2Quat(pert_quat, pert_axis, pert_angle);
            mju_rotVecQuat(vec_pert, vec_pert_scaled, pert_quat);
            printf("Internal perturbation of %f N delivered at time =  %f\n", pert_magn, d->time);
            printf("PD perturbation %f %f %f\n", c.PDpert[0], c.PDpert[1], c.PDpert[2]);
            for (int i = 0; i < 3; i ++) 
            {
                c.PDpert[i] = vec_pert[i];
            }
            //printf("pdvel %f, %f, %f\n", pdvel[0], pdvel[1], pdvel[2]);
            //printf("vec_pert %f, %f, %f\n", vec_pert[0], vec_pert[1], vec_pert[2]);
        }
    }
    else 
    {
        for (int i = 0; i < 6; i ++) {
            d->xfrc_applied[body_pert * 6 + i] = 0;
        }
        for (int i = 0; i < 3; i ++) {
            c.PDpert[i] = 0;
        }
        mju_copy(c.PDprev, c.PDmagn, 3);//PD vector at previous time step
    }
}

void vector2string(std::string result_string, std::string delimiter, double* vector, int vector_length)
{
    for (int i = 0; i < vector_length - 1; i++) 
    {
        result_string += (std::to_string(vector[i]) + delimiter);        
    }

    result_string += std::to_string(vector[vector_length - 1]);

    printf("this is a string, fool! %s\n", result_string.c_str() );

}
```
