## Supplementary material for "Visualising joint force-velocity properties in musculoskeletal models": Code&Data: README.md

# reaching-arm-model-CTR
 
This is a copy of the public reaching-arm-model-2.0 written by Tiina Murtola at the RVC, 2022,2023
It has been modified by Chris Richards at the RVC, 2023 to explore perturbations, FF vs FB control
and reach speed effects.  It will be made public upon publication.

All documentation for usage is in /docs

References:
[1]  Murtola, T., & Richards, C. (2023). The impact of intrinsic muscle properties on simulated reaching performance. Computer Methods in Biomechanics and Biomedical Engineering, 26(7), 777-788.
