## Supplementary figures and images for "Visualising joint force-velocity properties in musculoskeletal models"

### A03_FB_P00_f0_b.png

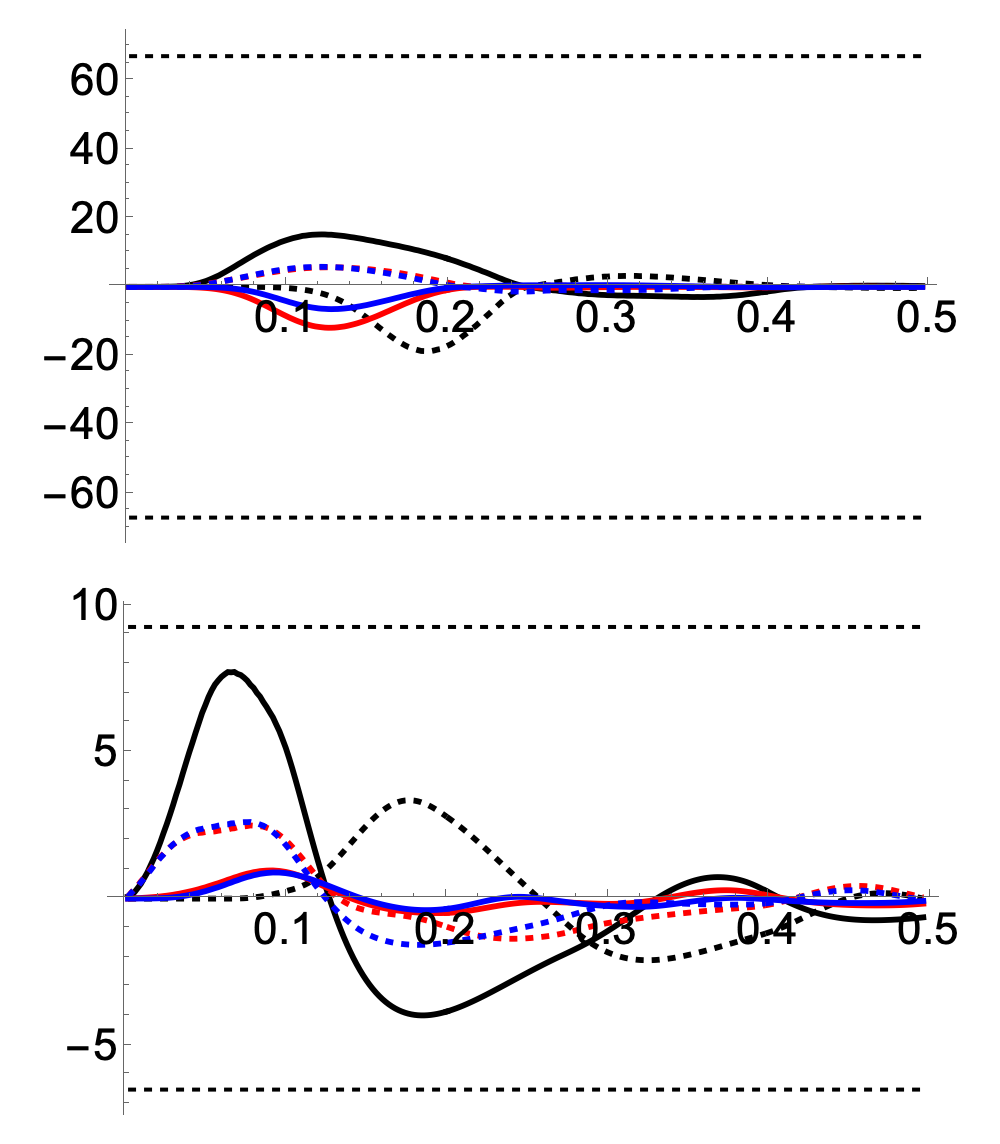
